# A piRNA-lncRNA regulatory network initiates responder and trailer piRNA formation during mosquito embryonic development

**DOI:** 10.1101/2020.03.23.003038

**Authors:** Valerie Betting, Joep Joosten, Rebecca Halbach, Melissa Thaler, Pascal Miesen, Ronald P. Van Rij

## Abstract

PIWI-interacting (pi)RNAs are small silencing RNAs that are crucial for the defense against transposable elements in germline tissues of animals. In *Aedes aegypti* mosquitoes, the piRNA pathway also contributes to gene regulation in somatic tissues, illustrating additional roles for piRNAs and PIWI proteins besides transposon repression. Here, we identify a highly abundant endogenous piRNA (propiR1) that associates with both Piwi4 and Piwi5. PropiR1-mediated target silencing requires base pairing in the seed region with supplemental base pairing at the piRNA 3’ end. Yet, propiR1 represses a limited set of targets, among which the lncRNA *AAEL027353* (*lnc027353*). Slicing of *lnc027353* initiates production of responder and trailer piRNAs from the cleavage fragment. Expression of propiR1 commences early during embryonic development and mediates degradation of maternally provided *lnc027353*. Both propiR1 and its lncRNA target are conserved in the closely related *Aedes albopictus* mosquito, underscoring the importance of this regulatory network for mosquito development.

## INTRODUCTION

PIWI proteins are a subfamily of the Argonaute protein family that associate with a specific class of 25-30 nt small non-coding RNAs to form RNA-induced silencing complexes (RISCs). Akin to small interfering (si)RNAs and micro (mi)RNAs, PIWI-interacting (pi)RNAs guide RISC to cognate RNAs through Watson-Crick base pairing to induce target silencing (Kobayashi & Tomari 2016). The piRNA pathway is mostly known for its role in silencing transposons to preserve genome integrity in the germline (Czech & Hannon 2016, Ozata et al. 2019). In *Drosophila*, the majority of piRNAs are produced from piRNA clusters, genomic regions that are rich in transposon remnants (Brennecke et al. 2007). piRNA cluster transcripts are transported to the cytoplasm, loaded onto the PIWI proteins Aubergine (Aub) and Piwi, and processed into mature piRNAs (Czech & Hannon 2016, Ozata et al. 2019). Whereas Piwi-piRNA complexes move to the nucleus to guide the deposition of repressive histone marks at transposon loci, piRNA-loaded Aub cleaves target RNAs in the cytoplasm through a mechanism termed slicing, thus repressing transposable elements at the post-transcriptional level (Gunawardane et al. 2007).

Efficiency of piRNA-mediated silencing relies on processes that amplify and diversify the pool of piRNAs. The 3’ cleavage fragments resulting from Aub-mediated slicing are bound by Ago3 and further processed into a responder piRNAs. In turn, Ago3-associated responder piRNAs cleave piRNA cluster transcripts, which are processed into new Aub-associated piRNAs, completing the so-called ping-pong amplification cycle. The preference of Aub to bind piRNAs with a 5’ terminal uridine, combined with the fact that PIWI proteins slice their targets between nucleotides 10 and 11 of their guide, gives rise to the 1U/10A signature that is characteristic of piRNA production through the ping-pong amplification loop (Gunawardane et al. 2007). The 3’ ends of piRNAs are defined by the endonuclease Zucchini (Zuc) in complex with accessory proteins at the mitochondrial membrane. In addition, the same complex processes the downstream 3’ fragment of the piRNA precursor into a series of trailer piRNAs, which predominantly associate with Piwi and direct transcriptional silencing of transposons in the nucleus (Han et al. 2015, Mohn et al. 2015).

Although the piRNA pathway is mostly studied in fruit flies, they represent an atypical case within the arthropod phylum, considering that their piRNA expression is restricted to germline tissues. In most arthropods, including *Aedes* mosquitoes, PIWI proteins and piRNAs are expressed in both germline and somatic tissues (Lewis et al. 2018). Moreover, the PIWI gene family has undergone expansion to seven members in *Aedes aegypti*, compared to three in *Drosophila* (Campbell et al. 2008, Lewis et al. 2016), suggesting that the pathway has undergone functional diversification in mosquitoes. Indeed, *Aedes* mosquitoes produce piRNAs from a diverse set of substrates and members of the PIWI family show specificity with regard to their piRNA repertoire. For instance, Piwi5 is required for the production of piRNAs derived from transposons, cytoplasmic RNA viruses, and endogenous viral elements. In contrast, Piwi4 is highly enriched for two piRNAs (tapiR1 and −2) derived from an evolutionary conserved satellite repeat locus, but depleted for transposon- or virus-derived piRNAs (Halbach et al. 2020, Miesen et al. 2015, Palatini et al. 2017, Suzuki et al. 2017, Whitfield et al. 2017). Additionally, Piwi5 and Ago3 are the core proteins in the ping-pong amplification loop that produces piRNAs from viral, transposon and messenger RNA substrates (Arensburger et al. 2011, Girardi et al. 2017, Joosten et al. 2018, Miesen et al. 2015).

The observation that *Ae. aegypti* produces abundant piRNAs mapping to protein-coding genes strongly suggests that piRNA function in *Aedes* mosquitoes extends to gene regulation (Arensburger et al. 2011, Girardi et al. 2017, Miesen et al. 2015). Indeed, we recently demonstrated that the Piwi4-associated piRNA tapiR1 regulates the expression of both protein-coding and non-coding RNAs during embryogenesis (Halbach et al. 2020). Early embryonic development of animals is driven by maternally provided mRNAs until maternal-to-zygotic transition (MZT), during which the zygotic genome is activated and maternal mRNAs are degraded (Tadros & Lipshitz 2009). Degradation of maternal mRNAs relies, at least in part, on maternally deposited piRNAs in fruit flies (Barckmann et al. 2015, Rouget et al. 2010), and on the zygotic piRNA tapiR1 in *Aedes* mosquitoes. As a consequence, selective inhibition of tapiR1 disrupts embryonic development, illustrating the importance of piRNA-mediated gene regulation during development (Halbach et al. 2020).

To further dissect the gene regulatory potential of individual endogenous piRNAs in *Ae. aegypti*, we describe here a highly abundant, Piwi4- and Piwi5-associated piRNA that we named propiR1. We found that this piRNA represses a limited set of targets, among which the long non-coding RNA (lncRNA) *AAEL027353*. propiR1 expression commences during the first hours of embryonic development to direct the degradation of *lnc027353*, suggesting that this regulatory network is important for embryonic development. Our data highlight the importance of regulatory networks that are governed by individual piRNAs to control various biological processes.

## RESULTS

### A host-derived piRNA associates with Piwi4 and Piwi5

To evaluate the gene regulatory potential of the *Ae. aegypti* piRNA pathway, we inspected small RNA deep-sequencing data from *Ae. aegypti* derived Aag2 cells to find highly abundant piRNAs. We focused on Piwi5-associated piRNAs, as Piwi5 is initially loaded with genome-encoded piRNAs and engages in efficient feed-forward piRNA amplification through the ping-pong loop (Joosten et al. 2018, Miesen et al. 2015). We selected the 25 most abundant Piwi5-associated piRNAs in Aag2 cells using previously published small RNA data from PIWI immunoprecipitations (IP) (Miesen et al. 2015). Strikingly, the Piwi5-associated piRNA with the highest expression in Aag2 cells did not map to transposon sequences (Fig. 1A).

**Figure 1.**
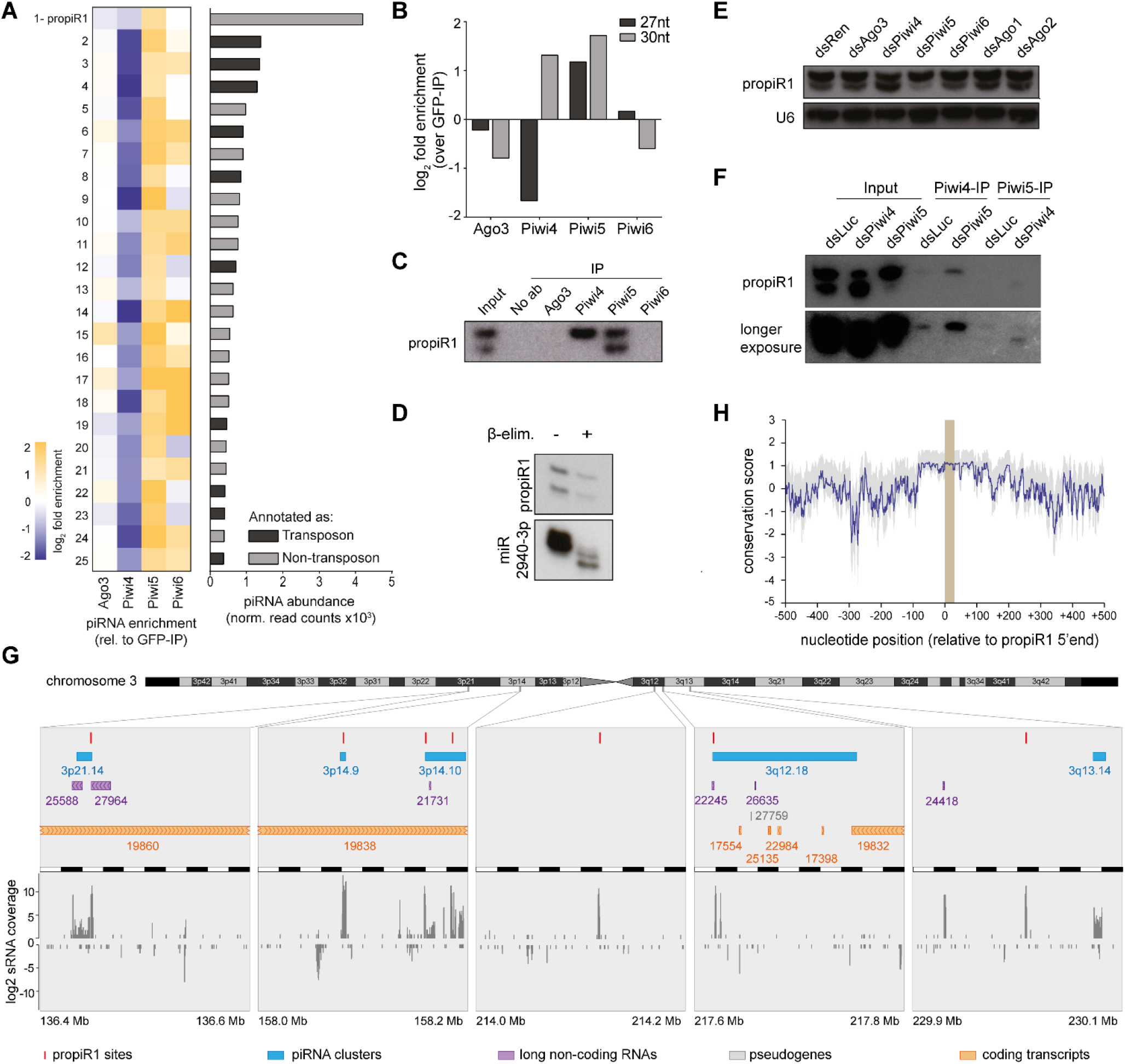
The endogenous piRNA propiR1 has two isoforms that differentially associate with Piwi4 and Piwi5. **A)** Ranked list of the top-25 most abundant piRNAs in Piwi5 immunoprecipitation (IP) in Aag2 cells. The heatmap (left) represents piRNA-enrichment in different PIWI protein IP libraries; the bar chart (right) shows piRNA abundance in Aag2 cells. Dark gray bars indicate transposon-derived piRNAs; light gray bars indicate piRNAs not mapping to annotated transposons. **B)** Enrichment of the two most abundant propiR1 isoforms (27 and 30 nt) in small RNA sequencing libraries generated from endogenous PIWI protein IPs. **C)** Northern blot analysis of propiR1 enrichment in PIWI protein IPs using antibodies targeting endogenous PIWI proteins from Aag2 cell lysates. An IP in which the antibody was omitted serves as control (No ab). **D)** Northern blot analysis of propiR1 and miR-2940-3p in RNA samples subjected to β-elimination as indicated. The blot was reused from Halbach et al. 2020 and re-probed for propiR1. The miR2940-3p panel is shown here again as a control, reproduced from Fig. 1B in Halbach et al. 2020. “A satellite repeat-derived piRNA controls embryonic development of Aedes.” *Nature*, published 2020 by Springer Nature. **E)** Northern blot analysis of propiR1 in Aag2 cells transfected with dsRNA targeting the indicated genes, or *Renilla* luciferase (dsRen) as a control. U6 snRNA serves as loading control. Knockdown efficiencies are shown in Supplemental Fig. S1C. **F)** Northern blot analysis of propiR1 in Piwi4 IP material from Aag2 cells treated with *Piwi5* or control dsRNA (Luciferase, dsLuc), and in Piwi5 IP material treated with *Piwi4* and control dsRNA. rRNA serves as loading control. **G)** Schematic representation of propiR1 sites on chromosome 3 of *Ae. aegypti* and their piRNA coverage. propiR1 sites are indicated in red (top row). Protein-coding and non-coding genes, pseudogenes and piRNA clusters (as annotated in VectorBase) in a 200 kb window flanking the propiR1 piRNA sites are color-coded as indicated. piRNA clusters (in blue) were named according to chromosomal location, as described in Crava et al. 2021. VectorBase gene identifiers (without the *AAEL0* species prefix) were used to refer to transcripts. Small RNA (19-32 nt) coverage in Aag2 cells is indicated relative to 10^6^ miRNAs. **H)** Sequence conservation between the seven propiR1 sites (marked as a gray box) and 500 bp flanking regions. The conservation scores (solid purple line) and 50% confidence intervals (gray shading) are indicated for each nucleotide position.

Upon closer inspection, we found that this piRNA was expressed as two distinct isoforms of 27 and 30 nt in size (Supplemental Fig. S1A). Small RNA deep sequencing as well as northern blotting revealed that both isoforms were enriched in Piwi5 IP, yet, the 30 nt isoform specifically interacted with Piwi4 (Fig. 1B-C, Supplemental Fig. S1B). This isoform specific Piwi4 enrichment was missed in the initial analysis, as we did not examine piRNA enrichment for the distinct sizes separately. Because of this dual PIWI protein association pattern, we named this piRNA promiscuous piRNA 1 (propiR1).

After loading onto a PIWI protein, the 3’ terminal ribose of piRNAs are 2’-*O*-methylated by the *S*-adenosylmethionine-dependent methyltransferase Hen1 (Horwich et al. 2007, Saito et al. 2007, Tian et al. 2011). The presence of this modification can be assessed by sodium periodate oxidation followed by β-elimination, which removes the 3’ nucleoside of unmodified RNAs, resulting in increased electrophoretic mobility (Kawaoka et al. 2014). In contrast to miRNA miR2940-3p, neither of the propiR1 isoforms are affected by β-elimination (Fig. 1D), indicating that these piRNAs have 2’-*O*-methylated 3’ ends and are mature, PIWI protein associated piRNAs. Moreover, this result suggests that the 30 nt isoform is unlikely to be a precursor of the 27 nt isoform.

To investigate the involvement of different PIWI proteins in propiR1 biogenesis, we assessed the effect of RNAi-mediated knockdown of *Ago3* and *Piwi4-6* (Supplemental Fig. S1C), as these PIWI genes are abundantly expressed in both germline and somatic tissues, as well as in the somatic Aag2 cell line (Akbari et al. 2013, Joosten et al. 2018, Vodovar et al. 2012). In accordance with the differential association of the two propiR1 isoforms, *Piwi4* knockdown resulted in a mild reduction of the larger 30 nt isoform, while abundance of the 27 nt isoform was clearly reduced upon *Piwi5* knockdown (Fig. 1E). As expected, knockdown of *Ago1* and *Ago2*, components of the miRNA and siRNA pathway, respectively, did not affect biogenesis of either propiR1 isoform (Fig. 1E).

Interestingly, we found that *Piwi4* knockdown resulted in an increased abundance of the short propiR1 isoform, suggesting an interplay between Piwi4 and Piwi5 in propiR1 biogenesis. We propose that competition between Piwi4 and Piwi5 for a putative propiR1 precursor may account for the observed effects of PIWI knockdown. To support this hypothesis, we analyzed propiR1 association with Piwi4 upon *Piwi5* knockdown and vice versa. Indeed, we found that more of the 30 nt isoform is bound by Piwi4 upon *Piwi5* knockdown, compared to control knockdown. Conversely, in *Piwi4* knockdown conditions, we observed increased binding of the short propiR1 isoform to Piwi5 (Fig. 1F).

propiR1 mapped to seven locations on chromosome 3 (Fig. 1G), five of which are inside piRNA clusters (Crava et al. 2021) These seven sites displayed a high degree of sequence conservation covering the piRNA and a ~250 bp flanking sequence element (Fig. 1H). The sequences outside this element were more divergent, suggesting that the piRNA site was duplicated on the chromosome as part of a larger fragment, of which the ~250 nt sequence element was preferentially maintained throughout evolution.

### propiR1 is able to silence target RNAs in trans

Studies in *Ae. aegypti* and *C. elegans* have shown that piRNAs silence mRNAs via miRNA-like recognition using a 5’ seed sequence (nt 2-7) with additional 3’ supplementary base pairing (Halbach et al. 2020, Shen et al. 2018). To evaluate propiR1 silencing potential and targeting requirements, we set up a reporter assay in which a single 30 nt target site for propiR1 was introduced into the 3’ UTR of firefly luciferase (FLuc) (Fig. 2A). This reporter thus contains a fully complementary target site for both the 27 nt and 30 nt isoforms. Compared to a reporter without target site, the reporter bearing a fully complementary propiR1 target site was silenced ~10-fold (Fig. 2B), indicating that propiR1 is capable of silencing target RNAs *in trans*.

**Figure 2.**
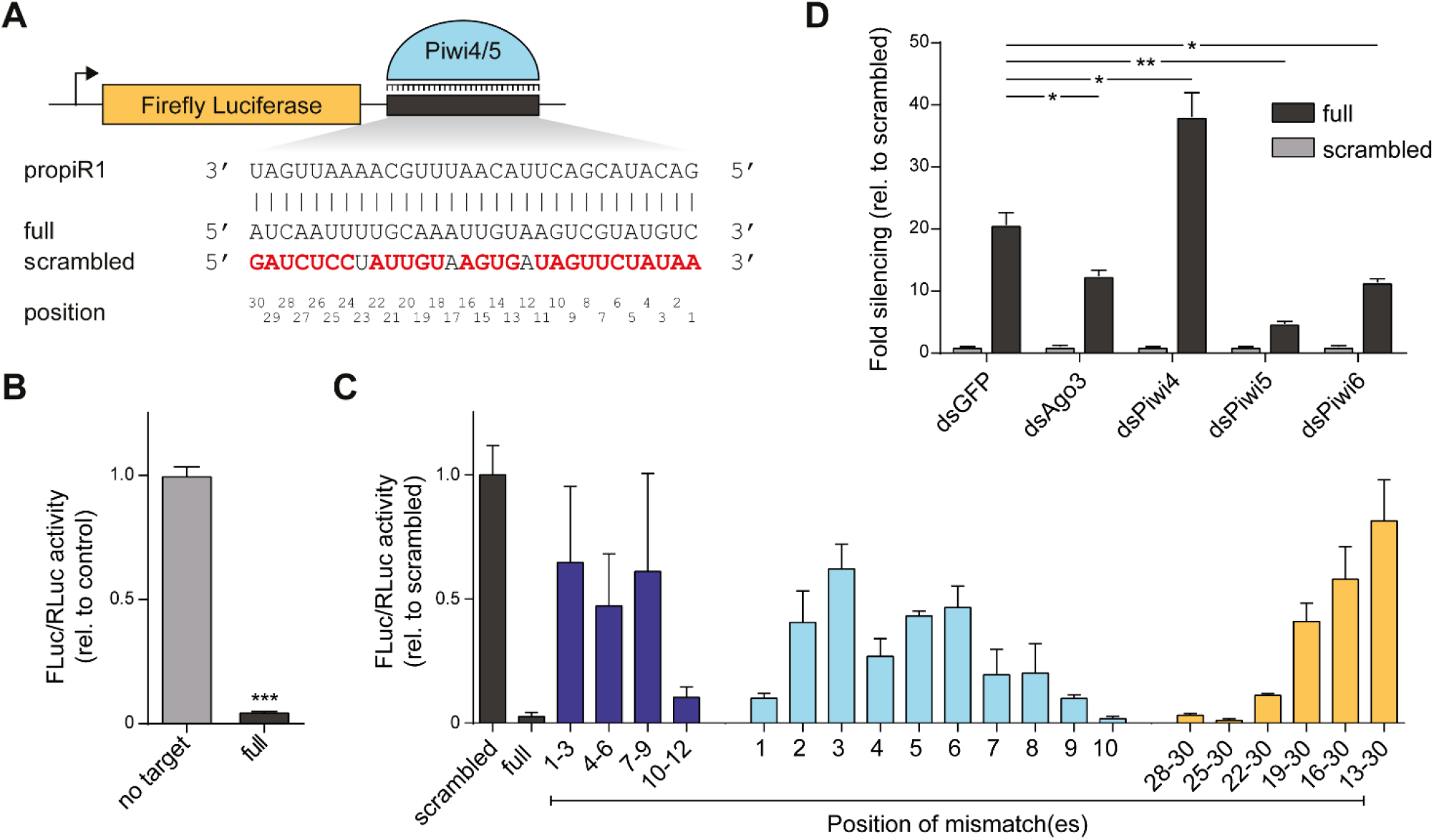
propiR1 has gene regulatory potential *in trans*. **A)** Schematic representation of the firefly luciferase (FLuc) reporters bearing a propiR1 target site or a control sequence in the 3’ UTR. Numbers correspond to the position of residues in the target RNA (t1-30) expected to base pair with propiR1 nucleotides 1-30 (from 5’ to 3’). Sequences of propiR1, a fully complementary target site, and the scrambled control (used in (C)) are shown. **B)** Luciferase assay using reporters bearing a fully complementary target site (full) or no target site. FLuc activity was normalized to a co-transfected *Renilla* luciferase (RLuc) reporter [same in (C) and (D)]. Bars represent mean +/- SD of a representative of five independent experiments, each performed with three biological replicates. *** *P* < 0.0005 (unpaired t-test). **C)** Luciferase assay of reporters with propiR1 target sites harboring three consecutive mismatches (dark blue) or single nt mismatches (light blue), or an increasing number of mismatches corresponding to the 3’ end of the piRNA (yellow). Numbers indicate positions in the target site that were mutated (t1-30, shown in (A)). In all cases, guanosine residues were changed to cytidine, adenosine to uridine, and *vice versa*. Bars depict mean +/- SD of a representative of three independent experiments, each performed with three biological replicates. **D)** Luciferase assay using reporters with either a fully complementary (full) or scrambled target site upon PIWI gene knockdown in Aag2 cells. dsRNA targeting GFP (dsGFP) was used as a control. Bars represent mean +/- SD of a representative of three independent experiments, each performed with three biological replicates. Asterisks denote statistically significant differences in fold silencing between dsGFP control and PIWI gene knockdowns (unpaired two tailed t-tests with Holm-Sidak correction; * *P* < 0.05, ** *P* < 0.005).

To assess propiR1 targeting requirements, we introduced a series of mismatches in the target sequence of the FLuc reporter (Supplemental Fig. S2A). As anticipated, three consecutive mismatches in the target sequence expected to base pair with nt 1-9 of propiR1 (t1-9) resulted in strong desilencing of the reporter (Fig. 2C, dark blue bars). Moreover, single nucleotide mismatches at positions t2-8 resulted in desilencing of the reporter (Fig. 2C, light blue bars). A mismatch at position t1 did not affect reporter silencing (Fig. 2C), which is in line with the observation that piRNA 5’ ends are tightly anchored in a binding pocket in the MID-domain of PIWI proteins (Matsumoto et al. 2016, Yamaguchi et al. 2020) and therefore do not contribute to target recognition. Increasing the number of mismatches in the target RNA corresponding to the piRNA 3’ end resulted in gradual desilencing of the reporter (Fig. 2C, yellow bars). Moreover, abolishing 3’ supplementary pairing altogether (mut t13-30) resulted in almost complete loss of silencing of the reporter. Guanosine and uridine residues can base pair and form so-called G:U wobbles. Such a G:U wobble was tolerated at t8, whereas a wobble at t5 resulted in similar desilencing as a mismatch at this position (Supplemental Fig. S2B). Altogether, our results establish nt 2-8 as the seed region of propiR1, at which base pairing is required for efficient targeting. Yet, seed-based target recognition alone is not sufficient for propiR1-mediated silencing and 3’ supplemental base pairing is essential for efficient silencing.

Argonaute proteins with slicing activity cleave their target RNAs between nucleotides 10 and 11 from the 5’ end of the guide RNA. Introducing a single mismatch at t10 or three consecutive mismatches that cover the putative slice site at t10-12 did not result in strong desilencing (Fig. 2C), suggesting either that a slicing-independent mechanism is responsible for propiR1-mediated silencing, or that base pairing at the slice site is dispensable for slicing.

To analyze PIWI protein dependency of reporter silencing, we performed a reporter assay upon knockdown of the four PIWI genes that are expressed in Aag2 cells. As control, we transfected cells with dsRNA targeting GFP (dsGFP), in which the reporter bearing the fully complementary propiR1 target site was silenced ~20-fold (Fig. 2D). Reporter silencing is reduced most prominently upon knockdown of *Piwi5* (~4.5-fold reduction compared to dsGFP; Fig. 2D). Yet, minor desilencing is also observed upon knockdown of *Ago3* (~1.7-fold reduction) and *Piwi6* (~1.8-fold reduction). Surprisingly, knockdown of *Piwi4* resulted in more efficient reporter silencing (~2-fold; Fig. 2D). We hypothesize that this increase may be the result of competition between Piwi4 and Piwi5 for a putative propiR1 precursor and increased levels of the short, Piwi5-associated propiR1 isoform upon *Piwi4* knockdown (Fig. 1E-F). We conclude that silencing of the reporter bearing a fully complementary target site is largely mediated by Piwi5 and that knockdown of *Piwi4* increases propiR1 loading onto Piwi5, leading to more robust Piwi5-mediated silencing.

### propiR1 regulates expression of lnc027353

To validate that reporter silencing was indeed mediated by propiR1, we designed a 2’-*O*-methylated antisense RNA oligonucleotide (AO) fully complementary to propiR1 that is expected to inhibit propiR1-mediated target silencing. Treatment of Aag2 cells with propiR1 AOs reduced the levels of both propiR1 isoforms (Supplemental Fig. S3A). In a luciferase assay in which cells were transfected with increasing amounts of AOs, reporter silencing was abolished, confirming that reporter silencing was indeed propiR1-dependent (Fig. 3A).

**Figure 3.**
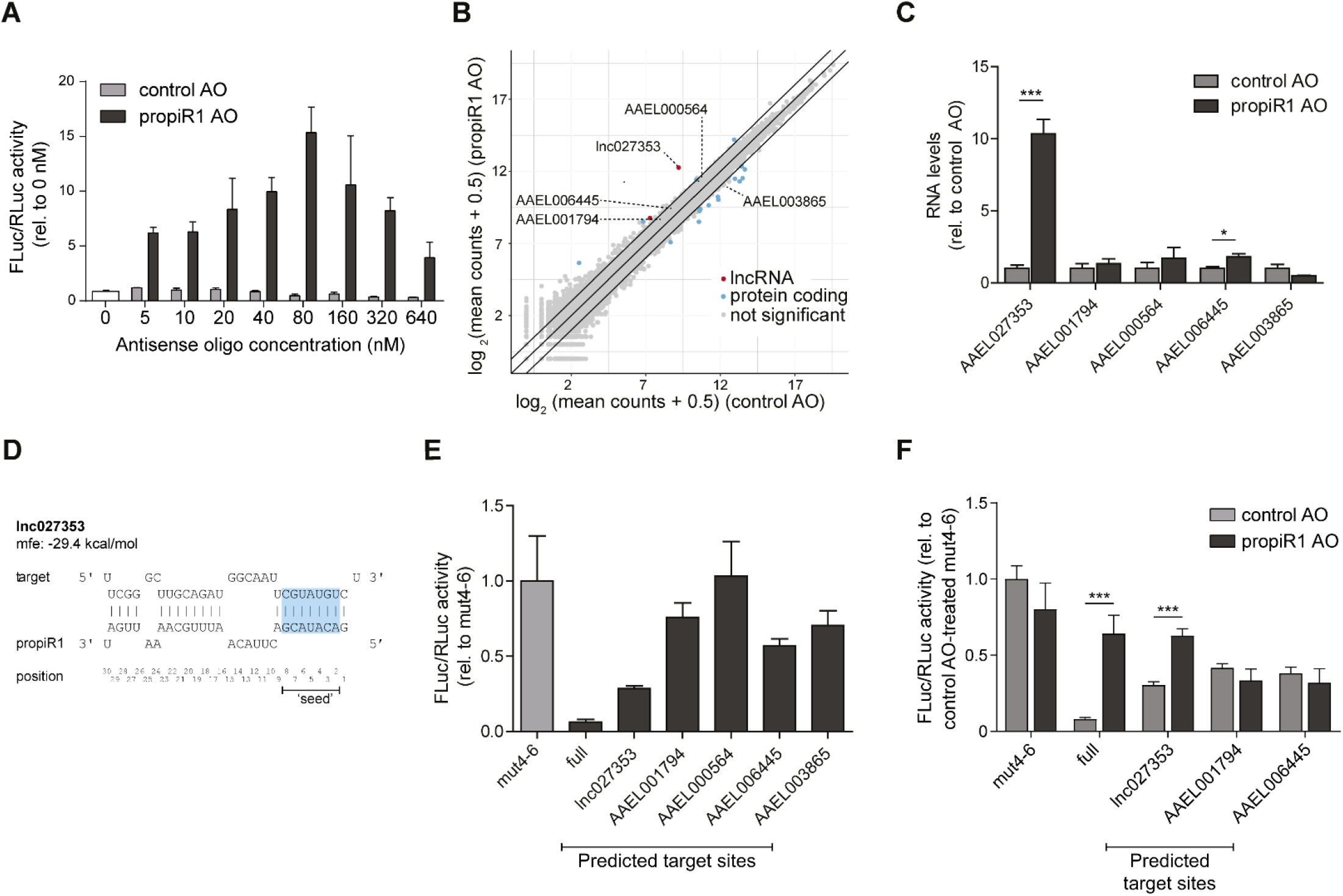
propiR1 regulates a limited set of cellular RNAs. **A)** Luciferase assay using a firefly luciferase (FLuc) reporter bearing a fully complementary propiR1 target site in the 3’UTR. Increasing amounts of 2’-*O*-methylated, antisense RNA oligonucleotides for propiR1 (propiR1 AO) or a control sequence (control AO) were co-transfected with the propiR1 reporter plasmid. FLuc activity was normalized to the activity of a co-transfected *Renilla* luciferase (RLuc) reporter [same in (E) and (F)]. Bars show the mean +/- SD of a representative of two independent experiments, each performed with three biological replicates. **B)** Log_2_ gene expression in Aag2 cells treated with propiR1 or control AOs. Mean RNA-seq counts of three biological replicates are shown (plus a pseudo-count of 0.5 to plot values of zero). Significance was tested at an FDR of 0.01 and log_2_ fold change of 0.5. Diagonal lines highlight 2-fold changes. Significant, differentially expressed lncRNAs and protein coding genes are indicated with red and blue dots, respectively; gray dots indicate genes that were not significantly affected by propiR1 AO treatment. **C)** RT-qPCR of genes bearing propiR1 target sites after treatment of Aag2 cells with propiR1 or control AOs for 24 hours. Shown are the mean +/- SD of a representative of three independent experiments, each performed with biological triplicates. Asterisks denote statistically significant differences in gene expression between propiR1 and control AO treated cells (unpaired two tailed t-tests with Holm-Sidak correction; *** *P* < 0.0005). **D)** Schematic representation of the predicted propiR1-*lnc027353* RNA duplex and its minimum free energy (mfe). Numbers indicate propiR1 piRNA positions (5’ to 3’) with the seed sequence (nt 2-8) in blue shading. **E)** Luciferase assay of reporters containing predicted propiR1 target sites from genes differentially expressed upon propiR1 AO treatment. Depicted are the mean +/- SD of a representative of three independent experiments, each with three biological replicates. Data are presented relative to the control reporter in which residues t4-6 were mutated (mut4-6). **F)** Luciferase assay of reporters containing predicted propiR1 target sites upon co-transfection with propiR1 or control AOs. Bars represent the mean +/- SD of four biological replicates. Data are presented relative to the mut4-6 control reporter treated with control AO. Asterisks denote statistically significant differences between control and propiR1 AO treatment (unpaired two tailed t-tests with Holm-Sidak correction; *** *P* < 0.0005).

To identify endogenous transcripts targeted by propiR1, we compared the transcriptome of Aag2 cells treated with propiR1 AOs to that of cells treated with control AOs using RNA-seq (Fig. 3B). propiR1 AO treatment resulted in a ~7.7-fold increase of a single lncRNA, *AAEL027353* (referred to as *lnc027353* throughout this study, *P* = 2.9E-64), while a handful of other lncRNAs and mRNAs were mildly affected as well (Fig. 3B, Supplemental Table S1). Furthermore, two transposon mRNAs were differentially expressed upon propiR1 AO treatment (Supplemental Fig. S3B, Supplemental Table S2). One of these (*Pao_Bel_Ele55*) contains a seed-like propiR1 target site (Supplemental Fig. S3C), which was a functional target site in a luciferase-based reporter assay (Supplemental Fig. S3D).

We used RNAhybrid (Rehmsmeier et al. 2004) to predict propiR1 target sites in affected genes wherein G:U wobbles at t2-7 were excluded (Supplemental Fig. S4A). Besides *lnc027353*, none of the significantly affected genes contained predicted propiR1 target sites. Hence, we selected the most strongly (albeit not significantly) affected genes bearing propiR1 target sites and evaluated their expression upon propiR1 AO treatment using RT-qPCR (Fig. 3C). While most changes in the expression of selected genes were mild and not significant, *lnc027353* expression was strongly increased (~10.3-fold) upon propiR1 AO treatment (Fig. 3C), akin to what we found using RNA-seq (Fig. 3B). Additionally, we analyzed *AAEL003865*, a gene that was downregulated in propiR1 AO-treated cells in RNA-seq (~2.1-fold) and were able to confirm this reduction (Fig. 3C), further validating our RNA-seq results.

To evaluate whether the changes in gene expression were due to direct propiR1-mediated targeting of these transcripts, we introduced the predicted target sites into the 3’ UTR of FLuc reporters (Fig. 3D, Supplemental Fig. S4A). Four out of five of the predicted target sites induced reporter silencing to some extent, of which the reporter containing the *lnc027353* target site was silenced most efficiently (~3.5-fold; Fig. 3E). Luciferase activity of the reporter containing the *AAEL003685* target site was not increased (Fig. 3E), suggesting that the reduction in *AAEL003865* expression upon selective propiR1 inhibition is likely an indirect, yet PIWI gene dependent effect (Supplemental Fig. S5).

To test whether the reporters with predicted target sites are specifically silenced by propiR1, we co-transfected the FLuc reporters with propiR1 or control AOs. Only the reporter bearing the predicted target site from *lnc027353*, but not from *AAEL001794* and *AAEL006445*, was desilenced by propiR1 AO treatment (Fig. 3F). propiR1 binds *lnc027353* target RNA through fully complementary base pairing across the seed region (nt 2-9), followed by a bulge that includes position 10 and 11, supplemented with incomplete base pairing at the 3’ end (Fig. 3D). Among the tested target sites, the propiR1-*lnc027353* target duplex had the lowest minimum free energy (−29.4, kcal/mol, Fig. 3D, Supplemental Fig. S4A), suggesting that binding efficiency contributes to target silencing efficiency. Of note, although RNAhybrid did not predict adenine-uridine base pairing at position 11 and 12, another RNA folding algorithm (Bifold, (Reuter & Mathews 2010)) did so in one of three alternative predictions, resulting in a less prominent unpaired bulge in the central region of the piRNA-target duplex (Supplemental Fig. S4B). Together, these results indicate that propiR1 efficiently and directly silences *lnc027353* through a partially complementary target site.

### PropiR1-mediated lnc027353 regulation is dominated by Piwi4

To dissect the contribution of the propiR1-binding PIWI proteins to silencing of endogenous targets, we performed *Piwi4* and *Piwi5* single and double knockdowns. In line with our previous findings (Fig. 1E), knockdown of *Piwi4* resulted in lower levels of the 30 nt isoform and higher levels of the Piwi5-bound 27 nt isoform (Fig. 4A), whereas *Piwi5* knockdown lead to reduced levels of both propiR1-isoforms. Combined knockdown of *Piwi4* and *Piwi5* resulted in the most pronounced reduction of both propiR1 isoforms (Fig. 4A). We next performed luciferase assays using reporters bearing propiR1 target sites in the context of single and double knockdown of *Piwi4* and *Piwi5*. As shown previously (Fig. 2D), silencing of a reporter bearing a fully complementary target site is partially alleviated by *Piwi5* knockdown (~3.5-fold desilencing; Fig. 4B), whereas *Piwi4* knockdown increased reporter silencing (~2-fold; Fig. 4B). In contrast, a reporter bearing the endogenous target site from *lnc027353* was desilenced upon *Piwi4* knockdown (~2-fold, Fig. 4B, Supplemental Fig. S6), while *Piwi5* knockdown had no effect.

**Figure 4.**
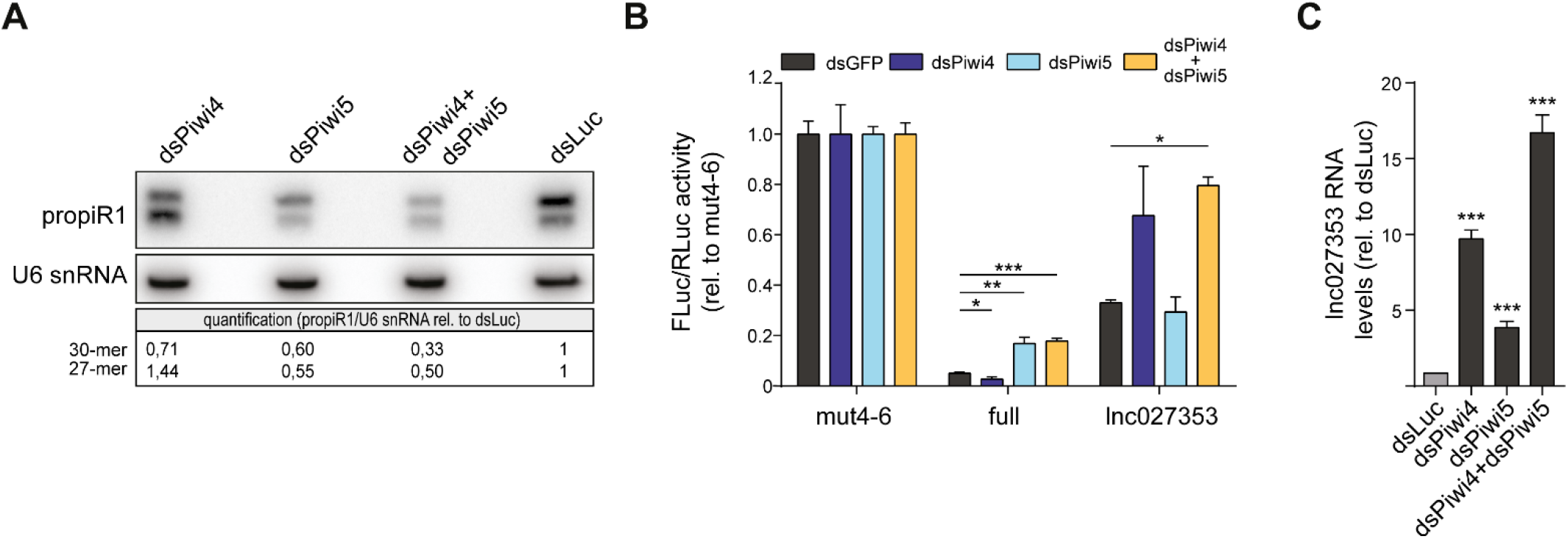
Piwi4 and Piwi5 adhere to different targeting rules. **A)** Northern blot analysis comparing the levels of the two propiR1 isoforms in *Piwi4* and *Piwi5* single and double knockdown (dsPiwi4, dsPiwi5) to a control knockdown (dsRNA targeting Luciferase, dsLuc). U6 snRNA serves as a loading control. The numbers below the blot are the quantified signal for each of the individual propiR1 isoforms, normalized to the U6 loading control, with the dsLuc control set to 1. **B)** Luciferase assay of reporters bearing a fully complementary propiR1 target (full), or the endogenous target site of *lnc027353* after single or double knockdown of *Piwi4* and *Piwi5*. A propiR1 target site in which residues t4-6 were mutated (mut4-6) serves as control. Luciferase activity was normalized to the mut4-6 control treated with the same dsRNA. Shown is a representative of four independent experiments, each performed with three biological replicates (the other experiments are shown in Supplemental Fig. S6). Bars represent mean +/- SD. Asterisks indicate statistically significant differences of the indicated reporters upon PIWI-gene knockdown compared to dsGFP-treatment (unpaired two tailed t-tests with Holm-Sidak correction; * *P* < 0.05, ** *P* < 0.005, *** *P* < 0.0005). **C)** Relative expression of the propiR1 target gene *lnc027353* upon *Piwi4* and *Piwi5* single and double knockdown in Aag2 cells, measured by RT-qPCR. Luc knockdown serves as a control and was used in single knockdowns to equalize the total amount of dsRNA per condition. Data represent the mean +/- SD of three biological replicates. Asterisks indicate statistically significant differences in *lnc027353* expression compared to dsLuc (unpaired two tailed t-tests with Holm-Sidak correction; *** *P* < 0.0005).

We next evaluated the effects of PIWI gene knockdown on expression of the endogenous propiR1 target *lnc027353*. *Piwi4* and *Piwi5* single knockdown resulted in a ~10-fold and ~4-fold increase in *lnc027353* expression, respectively (Fig. 4C). Knockdown of both *Piwi4* and *Piwi5* together resulted in a further increase of *lnc027353* expression (~17-fold, Fig. 4C), suggesting that both PIWI proteins contribute to lncRNA target silencing. While *Piwi5* knockdown moderately increased *lnc027353* expression (Fig. 4C), luciferase activity of a reporter bearing the *lnc027353* target site was not consistently affected by *Piwi5* knockdown (Fig. 4B, Supplemental Fig. S6). We excluded the possibility that other Piwi5-associated piRNAs are responsible for *lnc027353* degradation by testing the silencing efficiency of the nine most abundant non-propiR1 piRNAs in our FLuc reporter system (Supplemental Fig. S7). The reason for the inconsistency regarding the *Piwi5* knockdown effect on *lnc027353* reporter silencing and endogenous lncRNA expression remains unclear, but likely relates to the different readouts or cellular contexts in which these assays were performed. In the luciferase assay, the target site is studied upon transgene expression from a strong metallothionein promotor, whereas the RT-qPCR assesses endogenous lncRNA expression. Moreover, in contrast to the endogenous target RNA, the luciferase construct contains the 30 nt target site in a non-native sequence context. Irrespective, our data indicate that Piwi4 has a dominant role over Piwi5 in silencing the natural lncRNA target.

### propiR1 targeting initiates production of lnc027353-derived responder and trailer piRNAs

Target RNA slicing by PIWI proteins may trigger the production of responder and trailer piRNAs through ping-pong amplification and phased piRNA biogenesis, respectively (Gunawardane et al. 2007, Han et al. 2015, Mohn et al. 2015). Upon close inspection of small RNAs mapping to *lnc027353*, we found that the lncRNA transcript is indeed processed into responder and trailer piRNAs (Fig. 5A). Despite the fact that propiR1 and the *lnc027353* target site are not predicted to base pair at position t10 and t11 (Fig. 3D), we found ample production of responder piRNAs with a 5’-5’ offset of 10 nt respective to propiR1 (Fig. 5A). Furthermore, we detected low levels of trailer piRNAs generated from the *lnc027353* transcript, downstream of the propiR1 target site. These data suggest that propiR1 guides slicing of the lncRNA to initiate responder and trailer piRNA production.

**Figure 5.**
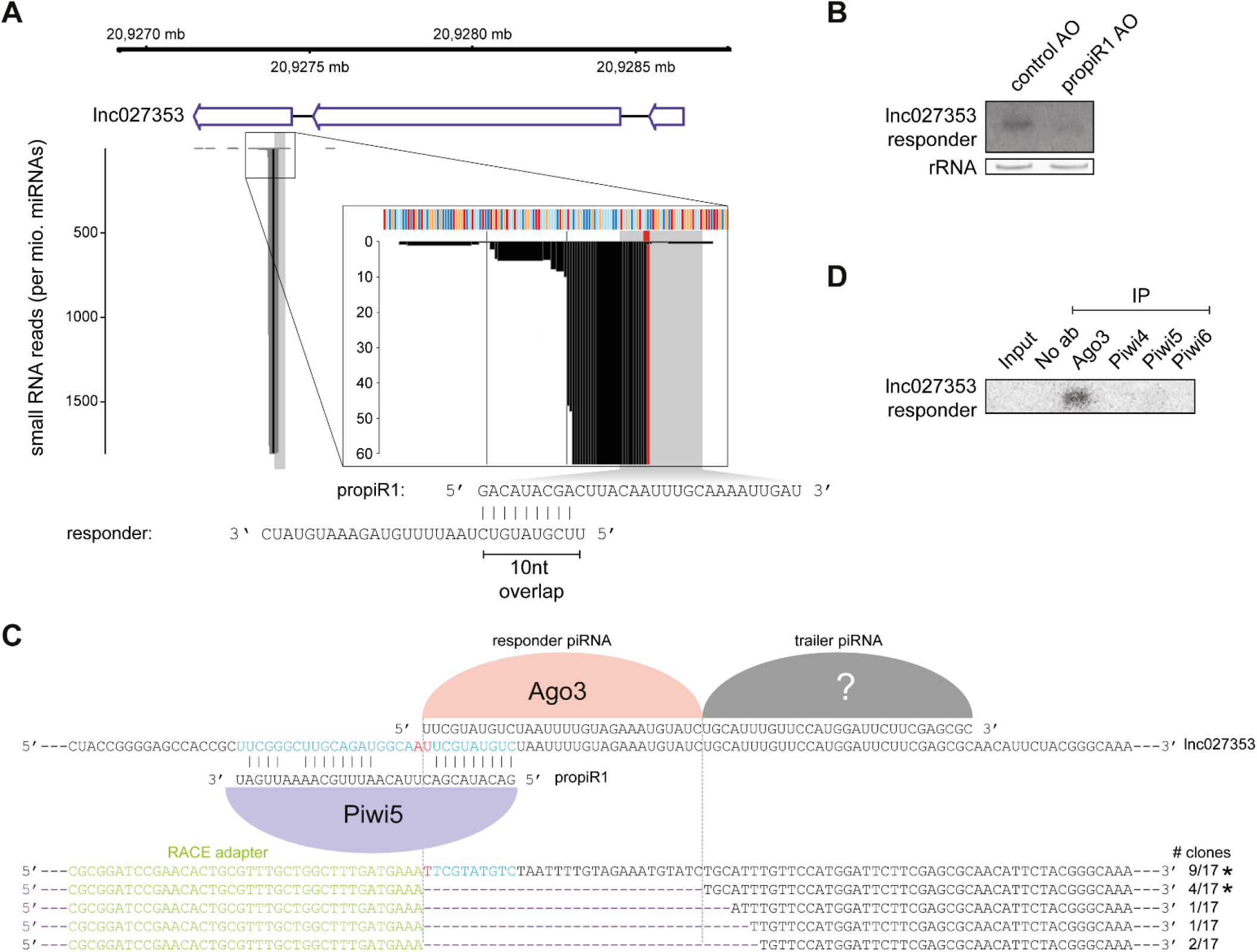
propiR1 targeting of *lnc027353* initiates responder and trailer piRNA production. **A)** Genomic coordinates, annotation and small RNA coverage of the *lnc027353* locus (top), with the position of the propiR1 target site indicated with a gray box. A magnification on the x-axis and y-axis is shown in the inset. The putative slice site (t10/11) is highlighted in red. Black vertical lines indicate distances of 30 nt from the putative slice site. The genomic sequence of the locus is represented with colored boxes (red: G; orange: C; light blue: A; dark blue: T). The sequence of propiR1 and the *AAEL027353*-derived responder piRNA are shown at the bottom. **B)** Northern blot analysis of the *lnc027353*-derived responder piRNA in cells treated with propiR1 or control antisense oligonucleotides (AO). EtBr stained rRNA serves as a loading control. **C)** Sanger sequencing results of 5’ RACE products of *lnc027353*, including the number of clones obtained (bottom panel) and schematic representation of generated piRNAs bound to different PIWI proteins (top panel). The expected slicer product as well as a putative Zucchini cleavage product serving as a precursor for trailer piRNA production are indicated with dashed vertical lines. Asterisks indicate 5’ RACE products corresponding to the 5’ ends of piRNAs detected by small RNA sequencing. One clone harboring the RACE adapter ligated 77 nt downstream of the slice site is not depicted. Green: 5’ RACE adapter; blue: propiR1 target site; red: slice site (t10/11), black: *lnc027353* transcript. **D)** Northern blot analysis of the *lnc027353*-derived responder piRNA in RNA extracted from immunoprecipitations (IP) of the indicated PIWI proteins. No ab indicates a control IP in which the antibody was omitted. The blot was re-used from Fig. 1C and re-probed for the *lnc027353* responder piRNA.

Treating Aag2 cells with propiR1 AOs, which reduce the expression of both propiR1 isoforms (Fig. 4A), strongly reduced the levels of the *lnc027353*-derived responder piRNA (Fig. 5B), indicating that its production is directly dependent on propiR1-mediated targeting. Indeed, using 5’ RACE we detected the 3’ cleavage product that is generated by propiR1-mediated slicing of *lnc027353*. Moreover, we recovered several cleavage products that are likely generated by Zucchini during downstream trailer biogenesis, with the cleavage site most often recovered by 5’ RACE also being supported by small RNA sequencing (Fig. 5C). In *Ae. aegypti*, Piwi5 and Ago3 are the core proteins in the ping-pong amplification complex, while Piwi4 appears to regulate gene expression independent of slicing activity (Halbach et al. 2020, Joosten et al. 2018, Miesen et al. 2015). We therefore hypothesize that Piwi5 slices *lnc027353* RNA, which should result in responder piRNA loading onto Ago3. In agreement, the *lnc027353*-derived responder piRNA was detected in Ago3 IP material (Fig. 5D), suggesting that it is indeed produced through a Piwi5 and Ago3-mediated ping-pong amplification loop.

While mismatches between guide and target RNA, especially around the slice site, are considered to abolish target RNA slicing by PIWI proteins, studies in various model systems show that low-level slicing does occur even in the absence of full complementarity around the slice site (Joosten et al. 2020, Mohn et al. 2015, Reuter et al. 2011). In line with these findings, our data provide evidence that propiR1 directs slicing of lnc027353, despite imperfect base pairing at and around the slice site. Our data cannot be used to infer efficiency of Piwi5-mediated slicing of *lnc027353* and it is likely that Piwi4 mediates silencing in a slicer independent manner, akin to previous observations with Piwi4/tapiR1 (Halbach et al. 2020). We propose that, even though Piwi4/propiR1 is dominant in silencing *lnc027353*, Piwi5/propiR1 slices a fraction of *ln027353* RNAs to generate responder piRNAs.

### The propiR1-controlled regulatory network is evolutionarily conserved

*AAEL027353* is annotated in VectorBase as a ~1.3 kb lncRNA containing two introns (Fig. 6A). However, publicly available RNA sequencing data (Akbari et al. 2013) suggest that only a small fraction (~300 bp) is transcribed, which includes parts of exons 2 and 3 and contains the propiR1 target site (Fig. 6A). Using RT-qPCR, we validated expression of the exon2-3 junction, as well as splicing of the intervening intron; yet, in line with the RNA sequencing data, we were unable to detect the exon1-2 junction (Supplemental Fig. S8A). Strikingly, the transcribed region of *lnc027353* coincides with an area of conservation between *Ae. aegypti* and *Ae. albopictus* (Fig. 6A; Supplemental Fig. S8B). The orthologous region (hereafter: *Aalb_lnc027353*) is not annotated as a gene in VectorBase, but could easily be detected by RT-qPCR, indicating that it is transcribed. This prompted us to investigate whether the regulatory circuit of propiR1 and *lnc027353* is conserved in *Ae. albopictus*.

**Figure 6.**
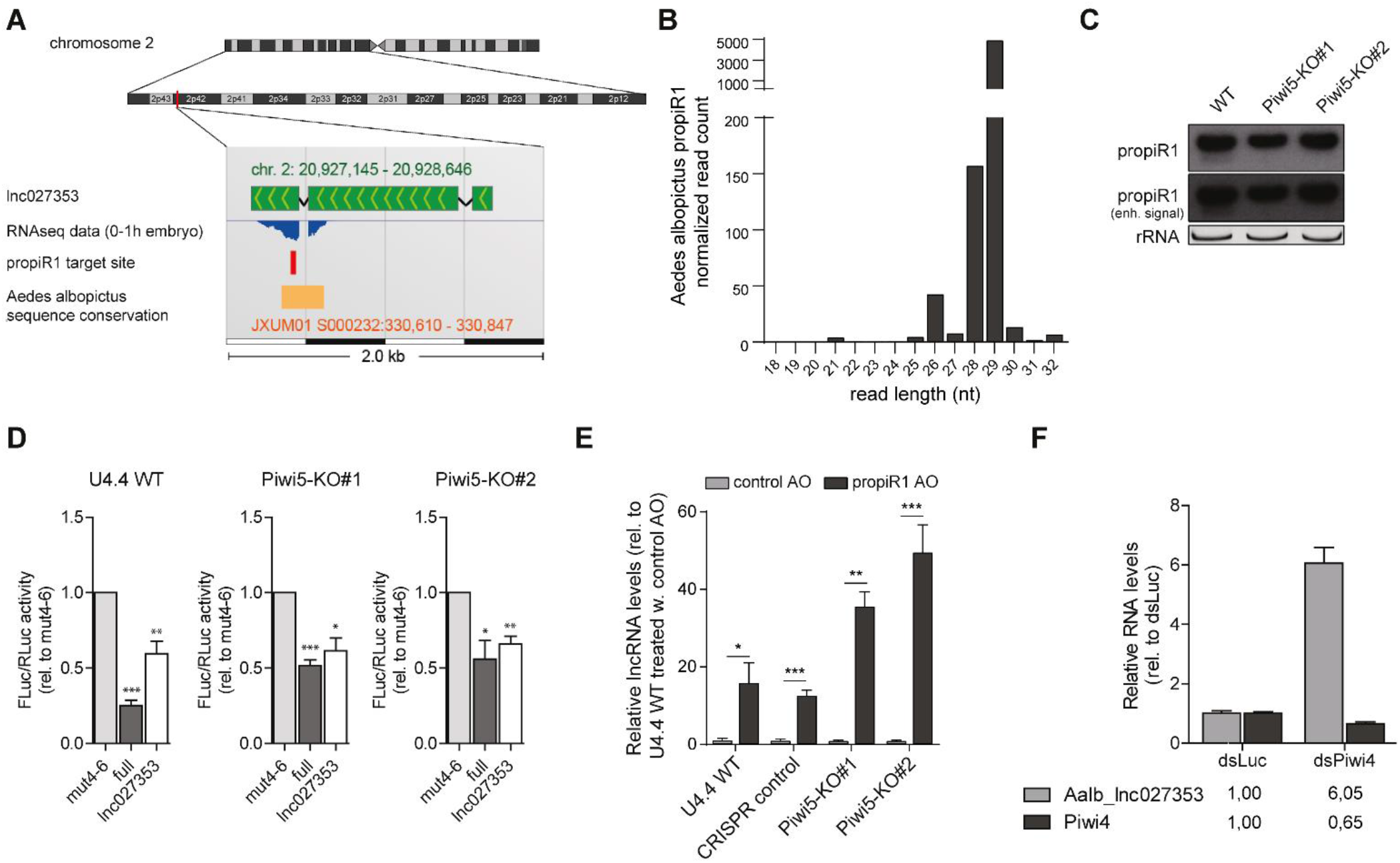
The propiR1- *lnc027353* regulatory network is evolutionary conserved. **A)** Schematic representation of the genomic location of *lnc027353* on *Ae. aegypti* chromosome 2 obtained from VectorBase. RNA-seq data from 0-1 h embryos (published under SRA accession SRS282183) are displayed using the APOLLO genome browser accessible through VectorBase. The position of the propiR1 target site (red) and an area with high degree of sequence conservation to *Ae. albopictus* (orange) are indicated. **B)** Size distribution of propiR1 isoforms in *Ae. albopictus* U4.4 cells. The read count is normalized to library size (per million reads). **C)** Northern blot analysis of propiR1 in wildtype (WT) and two *Piwi5* knockout clones (KO) of *Ae. albopictus-derived* U4.4 cells. The middle panel shows the propiR1 signal after the signal was enhanced to visualize the short propiR1 isoform. EtBr stained rRNA serves as loading control. **D)** Luciferase assay in WT and *Piwi5* KO U4.4 cells using reporters containing a control target site with mismatches at positions t4-6 (mut4-6), a fully complementary propiR1 target site (full) or the endogenous target site from *Ae. aegypti lnc027353*. Data were normalized to the activity of a co-transfected *Renilla* luciferase reporter (RLuc) and expression relative to mut4-6 is shown. Bars and whiskers represent the mean +/- standard error of the mean of three independent experiments (depicted in Supplemental Fig. S9), each performed with three biological replicates. Asterisks mark significant changes in luciferase activity between the different reporters within the same cell line (unpaired two tailed t-tests with Holm-Sidak correction; * *P* < 0.05, ** *P* < 0.005, *** *P* < 0.0005). **E)** Expression of the *Ae. albopictus lnc027353* ortholog in WT and *Piwi5* KO U4.4 cells, measured by RT-qPCR, after treatment with 300 nM propiR1 or control antisense oligonucleotides (AO) for 48 hours. CRISPR control is a clonal control cell line generated in parallel with the *Piwi5* KO cell lines. Bars and whiskers depict the mean +/- SD of three biological replicates. Asterisks mark significant changes in *lnc027353* expression upon transfection with control and propiR1 AOs within the same cell line (unpaired two tailed t-tests with Holm-Sidak correction; * *P* < 0.05, ** *P* < 0.005, *** *P* < 0.0005). **F)** Expression of the *Ae. albopictus lnc027353* ortholog and *Piwi4* in U4.4 cells upon *Piwi4* knockdown (dsPiwi4), measured by RT-qPCR. Data are normalized to cells treated with control dsRNA targeting luciferase (dsLuc). Bars and whiskers depict the mean +/- SD of three biological replicates.

propiR1 is highly abundant in *Ae. albopictus* U4.4 cells, however, unlike *Ae. aegypti* propiR1, *Ae. albopictus* propiR1 is almost exclusively expressed as a 29-mer, with a minor fraction being shorter (Fig. 6B). The shorter isoform was absent in *Piwi5* knockout (KO) U4.4 cells (Fig. 6C) [description and characterization of *Piwi5* KO cells in Taşköprü et al., in preparation], suggesting that, as in *Ae. aegypti*-derived Aag2 cells, the shorter isoform exclusively associates with Piwi5. In contrast, the longer 29 nt propiR1 isoform is abundantly expressed in *Piwi5* KO cells (Fig. 6C), indicating that its production does not require Piwi5, but is likely Piwi4 dependent.

The propiR1 target site is largely conserved in *Aalb*_*lnc027353* and is predicted to be targeted through seed-based and 3’ supplementary base pairing (Supplemental Fig. S8C, lower panel). Using the previously established luciferase assays, we found that the reporter bearing a target site fully complementary to *Ae. aegypti* propiR1 (which contains a single mismatch to *Ae. albopictus* propiR1; Supplemental Fig. S8C, upper panel) is consistently silenced in both wildtype (WT) and *Piwi5* KO U4.4 cells (Fig. 6D, Supplemental Fig. S9). Likewise, the reporter bearing the *lnc027353* target site is efficiently silenced in both WT and *Piwi5* KO cells (Fig. 6D, Supplemental Fig. S9). Thus, silencing of propiR1 target reporters is independent of Piwi5, and instead likely mediated by Piwi4 in *Ae. albopictus*.

To confirm propiR1-mediated regulation of the lncRNA in *Ae. albopictus* cells, we transfected propiR1 AOs into WT and *Piwi5* KO U4.4 cells. We found that expression of *Aalb_lnc027353* was increased ~16-fold in WT U4.4 cells and ~13-fold in a clonal CRISPR control U4.4 cell line upon propiR1 AO treatment (Fig. 6E), indicating strong propiR1-mediated regulation. In propiR1 AO treated *Piwi5* KO cells, lncRNA expression increased even more dramatically (~39–55-fold; Fig. 6E). These results indicate that *Aalb_lnc027353* is silenced in a Piwi5 independent manner. Indeed, knockdown of *Piwi4* resulted in strongly increased *Aalb_lnc027353* expression (~6-fold, Fig. 6F), indicating that *lnc027353* is silenced by Piwi4. For *Ae. aegypti*, we proposed that Piwi5-mediated slicing of *lnc027353* initiates responder piRNA production. In *Ae. albopictus*, however, barely any responder piRNAs are generated from *Aalb_lnc027353* in U4.4 cells (Supplemental Fig. S8D), suggesting the transcript is not efficiently sliced, likely due to the low expression of the Piwi5-associated short propiR1 isoform (Fig. 6B-C).

Together, these results indicate that like in *Ae. aegypti*, propiR1 regulates expression of *lnc027353* in *Ae. albopictus*. Interestingly, within the *Aalb_lnc027353* sequence, the propiR1 target site and the downstream sequence shows a higher degree of sequence similarity to *Ae. aegypti lnc027353* compared to the rest of the transcript (Supplemental Fig. S8B). Hence, the regulatory circuit between propiR1 and its lncRNA target is conserved between different *Aedes* species, suggesting its evolutionary importance.

### The propiR1-lnc027353 regulatory circuit is active in vivo

*Lnc027353* is expressed in all *Ae. aegypti* tissues including germline tissues (ovaries, testes, and sperm) (Supplemental Fig. S10A). Moreover, the transcript is abundant in early embryos (Supplemental Fig. S10B), which are transcriptionally silent, suggesting that it is maternally deposited. In contrast to its target, propiR1 is not maternally deposited, but its expression becomes detectable by northern blot at around 5 hours post oviposition (Fig. 7A). Initially, only the 30-nt isoform is expressed, but the 27-nt isoform becomes detectable 48 hours post oviposition (Fig. 7A).

**Figure 7.**
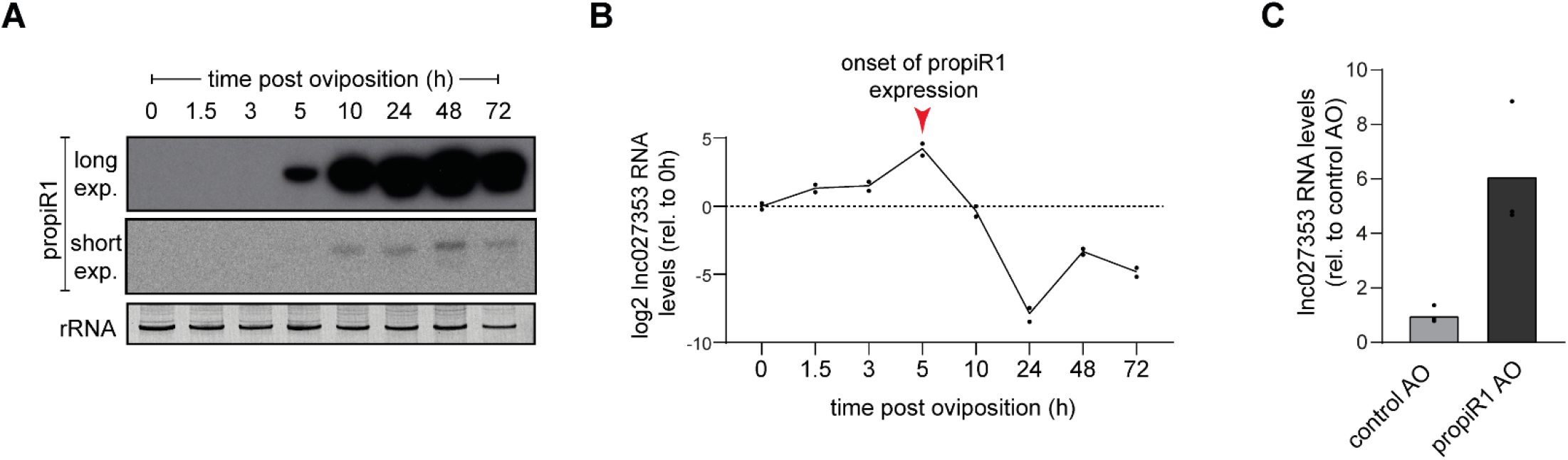
propiR1 regulates *lnc027353* expression during early development. **A)** Northern blot analysis of propiR1 in early embryonic development, with two different exposure times. Time indicates the age of the embryo in hours after a 30-minute egg laying period (oviposition). For each time point, RNA from 200-400 eggs was pooled. The northern blot, previously used in Halbach et al. 2020, was re-probed with a propiR1 probe. The EtBr stained rRNA panel is shown here again as loading control, reproduced from Fig. 3A in Halbach et al. 2020. “A satellite repeat-derived piRNA controls embryonic development of Aedes.” *Nature*, published 2020 by Springer Nature. **B)** *lnc027353* expression during early embryonic development, as measured by RT-qPCR. Line and dots represent mean and individual measurements of two technical replicates, respectively. Expression was normalized to the 0 h time point. The same RNA was used in (A) and (B). **C)** *lnc027353* expression in embryos injected with propiR1 or control antisense oligonucleotides (AO), as measured by RT-qPCR. Per condition, ~50 embryos were pooled. A representative of three independent experiments is shown. Bars and dots indicate the mean and individual measurement of technical triplicates, respectively.

As propiR1 expression is maintained in most tissues in the adult mosquito (Supplemental Fig. S10C), it may have diverse regulatory functions in adulthood. Yet, the dynamic regulation of propiR1 and its target *lnc027353* during embryogenesis prompted us to investigate the regulatory circuit in early development. Interestingly, we observed that the decrease of *lnc027353* expression coincided with onset of propiR1 expression (Fig. 7B, Supplemental Fig. S10B). To confirm that the reduction of *lnc027353* expression was propiR1 dependent, we injected *Ae. aegypti* embryos with propiR1 AOs. As hypothesized, this treatment resulted in a ~6-fold increase in *lnc027353* levels (Fig. 7C), indicating propiR1 is indeed responsible for silencing *lnc027353* expression during early embryonic development.

## DISCUSSION

In *Ae. aegypti*, the PIWI gene family has undergone expansion to seven members, whereas the genetic model organism *Drosophila melanogaster* only encodes three. This expansion, together with the fact that PIWI proteins are expressed in somatic as well as germline tissues suggests that the piRNA pathway has gained additional functions in *Aedes* mosquitoes, aside from the repression of transposons in the germline. It has been proposed that the *Ae. aegypti* piRNA pathway is involved in various processes including antiviral defense, regulation of gene expression, and degradation of maternally provided transcripts during the maternal to zygotic transition (Girardi et al. 2017, Halbach et al. 2020, Miesen et al. 2015, Morazzani et al. 2012, Schnettler et al. 2013, Tassetto et al. 2019, Vodovar et al. 2012).

To reveal further non-canonical functions of the piRNA pathway in *Ae. aegypti*, we identified highly abundant endogenous piRNAs not originating from transposable elements. We found that one of these piRNAs (propiR1) has strong silencing potential and set out to characterize its function. In *Ae. aegypti* Aag2 cells, propiR1 is expressed as two isoforms, which differentially associate with Piwi4 and Piwi5. propiR1-mediated target silencing requires Watson-Crick base pairing in the seed region plus additional supplemental pairing at the 3’ end. Given these lenient target sequence requirements, it is somewhat surprising that propiR1 strongly silences only a single non-transposon target in Aag2 cells: *lnc027353*. However, it should be noted that, given its ubiquitous expression during development and in various adult mosquito tissues, propiR1 may have additional *in vivo* targets that are not expressed in Aag2 cells. propiR1 mediates silencing of *lnc027353* during early embryonic development and propiR1-mediated cleavage results in the production of responder and trailer piRNAs from the cleavage fragment. Additionally, we show that the regulatory circuit of propiR1 and *lnc027353* is conserved in *Ae. albopictus*, which underscores its importance for mosquito development. Together, our results illustrate an intriguing interplay between two isoforms of a single piRNA, loaded onto two PIWI proteins, and a lncRNA during embryonic development of *Aedes* mosquitoes.

### Piwi4 and Piwi5 compete for a putative propiR1 precursor

It has been hypothesized that members of the expanded PIWI gene family have specialized to preferentially process different RNA substrates (Lewis et al. 2016, Miesen et al. 2015, Miesen et al. 2016). Apart from differences in the origin of RNA substrates that are processed into piRNAs, the spectrum of downstream mRNA targets might also be determined by differences in targeting requirements between PIWI proteins. The fact that propiR1 associates with two PIWI proteins (Piwi4 and Piwi5) allowed us to investigate variations in the targeting requirements of two PIWI proteins guided by the same small RNA.

We found that silencing of reporters with a fully complementary propiR1 target site is reduced upon *Piwi5* knockdown, demonstrating that silencing of such targets is *Piwi5*-dependent. In line with this, reporters bearing a fully complementary target site are silenced more efficiently in *Piwi4* knockdown, due to an increased availability of the propiR1 substrate to be processed into Piwi5-bound piRNAs. In contrast, *Piwi4* knockdown alleviated silencing of the *lnc027353* target gene as well as reporters containing the *lnc027353* target site, which contains mismatches at positions t10-15, t24-25 and t30. Altogether, these results indicate that silencing of a fully complementary propiR1-target site is achieved predominantly by Piwi5, whereas targeting of the imperfect target site in *lnc027353* results in Piwi4-mediated repression.

Piwi5 is known to have slicing activity, as is evident from its involvement in the ping-pong amplification loop (Joosten et al. 2018, Miesen et al. 2015). Slicing-mediated target silencing is expected to require more extensive base pairing compared to slicing-independent silencing mechanisms (Reuter et al. 2011) and therefore, it is expected that Piwi5-mediated silencing only allows a limited number of mismatches between piRNA and target RNA. Surprisingly, the production of responder and trailer piRNAs from the *lnc027353* cleavage fragment and the detection of slice products, illustrates that the transcript is at least in part silenced through slicing, despite the fact that the propiR1 target site in *lnc027353* contains mismatches at the putative slice site. The exact sequence requirements necessary to accommodate slicing and subsequent responder piRNA production remain hitherto unexplored, but it has been observed before that slicing may occur, albeit inefficiently, in the absence of complete base pairing between small RNA and target RNA in the seed region and the slice site (Joosten et al. 2020, Mohn et al. 2015, Reuter et al. 2011).

Since propiR1-mediated regulation of *lnc027353* mainly depends on Piwi4 rather than Piwi5 in both *Ae. aegypti* and *Ae. albopictus*, the majority of *lnc027353* transcripts is likely regulated via slicing-independent mechanisms. As previously observed for the Piwi4/tapiR1 complex (Halbach et al. 2020), Piwi4/propiR1-mediated targeting appears to be more ‘miRNA-like’ and likely requires the recruitment of accessory protein complexes to establish slicing-independent target silencing. Despite extensive investigation, the molecular machinery that is recruited to achieve Piwi4-mediated target silencing remains enigmatic.

As *Piwi4* and *Piwi5* have different spatiotemporal expression profiles (Akbari et al. 2013, Danet et al. 2019, Matthews et al. 2016, Wang et al. 2018), they may enforce propiR1-mediated regulation of various biological processes, thus expanding the regulatory potential of propiR1. It remains to be explored to what extent other piRNAs employ multiple PIWI-proteins in the regulation of host genes.

### The propiR1-lnc027353 regulatory circuit in embryonic development

The target of propiR1, *lnc027353*, is a maternally provided transcript that is degraded during the first hours of embryonic development. This reduction in *lnc027353* RNA levels coincides with an increase in propiR1 abundance and blocking propiR1 function with antisense oligonucleotides resulted in stabilization of the *lnc027353* transcript, confirming that propiR1 is indeed responsible for *lnc027353* degradation. While Piwi4 is maternally provided, *Piwi5* expression commences later in embryonic development, after zygotic genome activation (Akbari et al. 2013). This is in line with the appearance of the distinct propiR1 isoforms during embryonic development, since the Piwi4-associated 30 nt isoform is already detectable at 5 hours post oviposition, whereas the Piwi5-associated 27 nt isoform appears only after 48 hours. Hence, it is likely that *lnc027353* degradation is achieved by an accessory complex deposited by propiR1-guided Piwi4. This does not, however, rule out a role for Piwi5 in *lnc027353* regulation at later stages of embryonic development and in other life stages of the mosquito.

The importance of piRNAs during embryonic development is supported by the fact that piRNA levels peak during embryonic development (Brennecke et al. 2008, Liu et al. 2016), and by their involvement in the degradation of maternal transcripts during MZT in both fruit flies and *Aedes* mosquitoes (Barckmann et al. 2015, Halbach et al. 2020, Rouget et al. 2010). We hypothesize that lncRNA degradation induced by propiR1 in *Aedes* mosquitoes is required for proper embryonic development, however, the exact function of *lnc027353* remains to be explored.

### Evolutionary conservation of the network

We found that *Ae. albopictus* also expresses propiR1 and a putative *lnc027353* ortholog in which the sequence adjacent to the propiR1 target site is preferentially conserved. The high degree of sequence conservation of especially the target site suggests that this genomic region has important functions that have been maintained during evolution. Although not annotated as a repeat, propiR1 is encoded by seven and ten repetitive loci in *Ae. aegypti* and *Ae. albopictus*, respectively. Notably, propiR1 contains a single nucleotide deletion in *Ae. albopictus* compared to the *Ae. aegypti* sequence. Since all propiR1 loci in *Ae. albopictus* share this deletion, the most parsimonious interpretation is that the genome of the common ancestor of *Ae. aegypti* and *Ae. albopictus* contained a single propiR1 locus, which has duplicated independently in both species.

Our newly discovered regulatory circuit in which propiR1 targets and degrades a single lncRNA target illustrates an intriguing non-canonical role for the piRNA pathway. The conservation of propiR1 as well as its target site in the *lnc027353* ortholog in *Ae. albopictus*, together with its regulatory role in the early embryo suggests that the circuit is crucial for mosquito development. Moreover, it was recently proposed that piRNA-mediated regulatory networks such as the one described in our study also exist in *Anopheles* mosquitoes, suggesting that piRNAs may play a central role in mosquito gene regulation (Jensen et al. 2020). propiR1 is one of millions of different piRNA sequences (Arensburger et al. 2011), which may differ in their spatiotemporal expression patterns. Hence, the gene-regulatory potential of the *Aedes* piRNA pathway is likely to be highly complex and many interesting regulatory circuits remain to be explored.

## MATERIALS AND METHODS

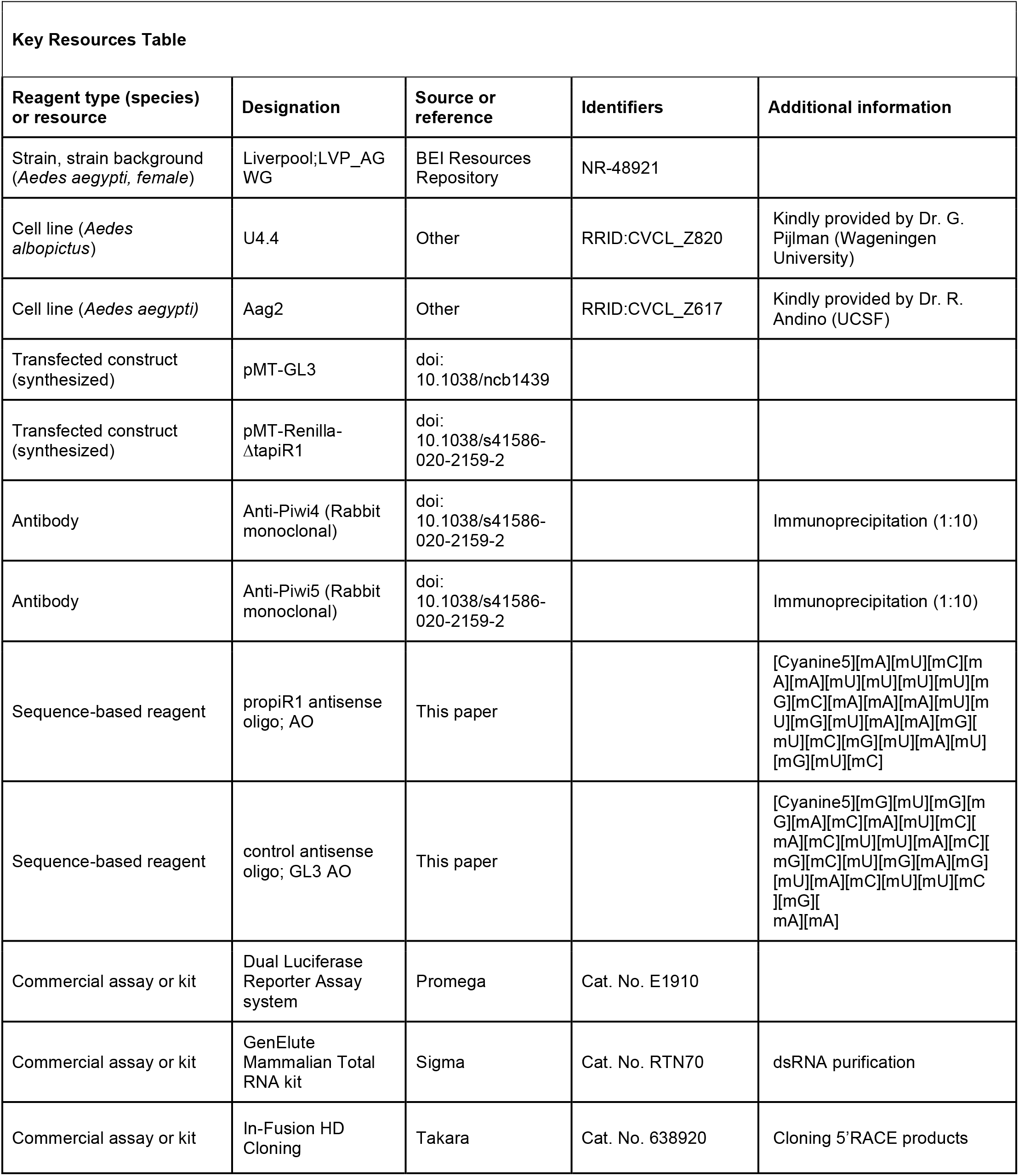
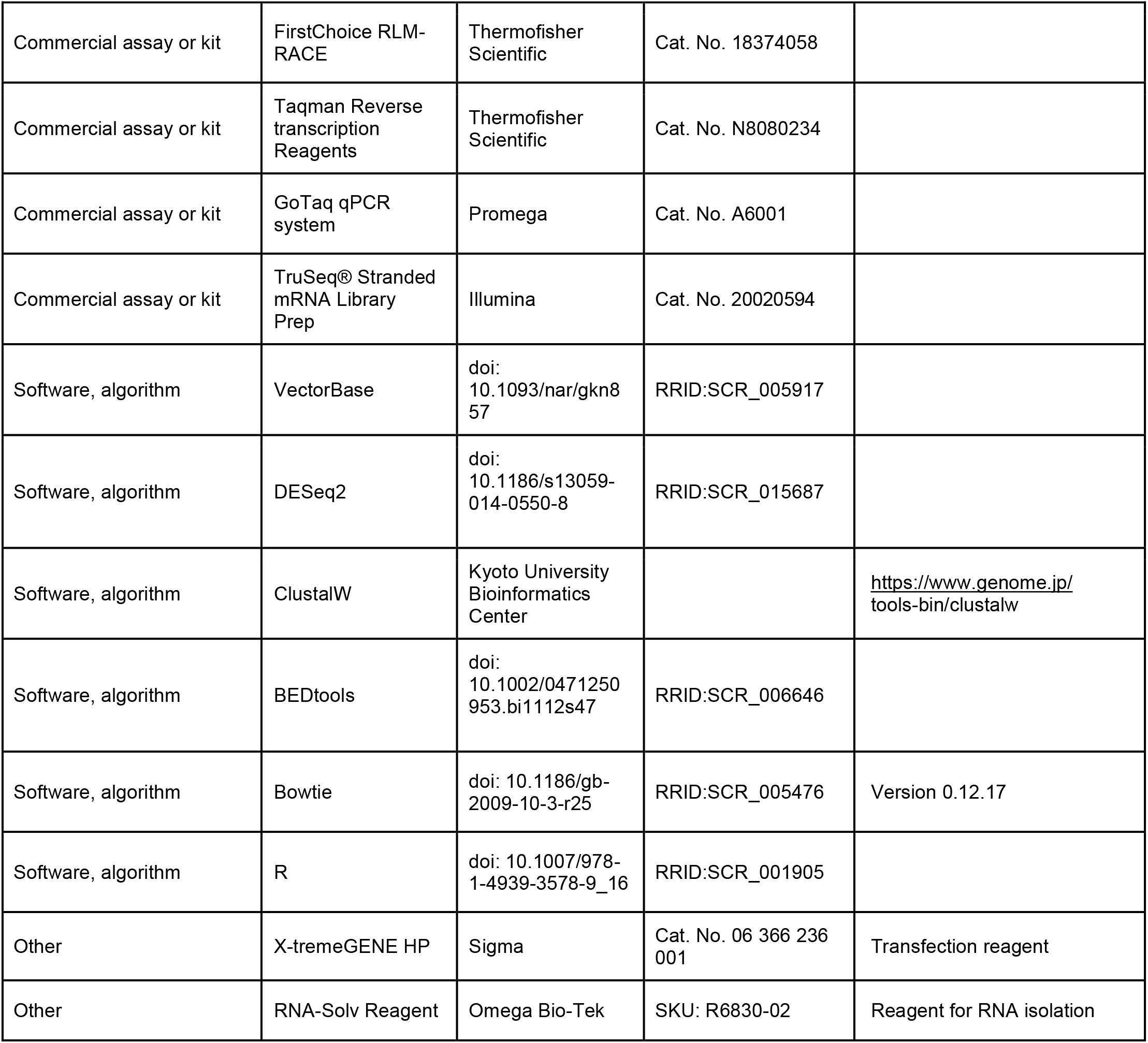

### Cell culture

*Aedes aegypti* Aag2 cells (RRID:CVCL_Z617, kindly provided by Raul Andino (UCSF)) and *Aedes albopictus* U4.4 cells (RRID:CVCL_Z820, kindly provided by Gorben Pijlman (Wageningen University)) were grown at 27°C in Leibovitz’s L-15 medium (Invitrogen) supplemented with 10% heat-inactivated fetal bovine serum (Gibco), 2% tryptose phosphate broth (Sigma), 50 U/mL Penicillin, 50 μg/mL Streptomycin (Gibco), and 1% Non-essential amino acids (Gibco). Cell lines were tested negative for mycoplasm. Cell lines were not authenticated, but (small)RNAseq data confirmed that they are *Ae. aegypti* and *Ae. albopictus* derived.

### Gene-knockdown using dsRNA

PCR products containing a T7 promoter sequence at both ends were generated and *in vitro* transcribed by T7 polymerase, heated to 80°C and gradually cooled down to form dsRNA. The dsRNA was purified using the GenElute Mammalian Total RNA kit (Sigma). Primers that were used to generate double stranded RNA are described in (Miesen et al. 2015). For knockdown experiments, cells were seeded at ~1 × 10^5^ cells/well in a 24-well plate and allowed to attach overnight. On the following day, 150 ng dsRNA/well was transfected into cells using 0.6 μL X-tremeGENE HP DNA Transfection Reagent (Sigma) according to the manufacturer’s instructions. Three hours later, medium was replaced by fresh, supplemented L15-medium. To improve knockdown efficiency, cells were transfected again in the same way 48 hours after the initial knockdown. 48 hours after the second transfection, cells were harvested in 1 mL RNA-Solv Reagent (Omega Bio-Tek) for RNA isolation.

### RNA isolation

Cells were homogenized in 1 mL RNA-Solv Reagent and subsequently, 200 μL of chloroform was added. After harsh vortexing and centrifugation, the aqueous phase was collected, and RNA was precipitated in 1.5 volumes of isopropanol for ~30 minutes on ice. RNA was pelleted by centrifugation at ~18,000 × g for 30 minutes at 4°C. Subsequently, pellets were washed 2-3 times in 85% ethanol, air-dried, and dissolved in nuclease free water.

### Periodate treatment and β-elimination

3’ modification of propiR1 was analyzed by re-probing a northern blot previously used in Halbach et al. 2020 with a propiR1 probe. For comparative reasons, the miR2940-3p blot is shown here again.

### Small RNA Northern blot

5-10 μg RNA was diluted in 2x RNA loading dye (New England Biolabs) and size separated on 7 M Urea/15% polyacrylamide/0.5×TBE gels by gel electrophoresis. RNA was transferred to Hybond Nx nylon membranes (Amersham) and crosslinked using EDC (1-ethyl-3-(3-dimethylaminopropyl)carbodiimide hydrochloride) (Sigma) (Pall & Hamilton 2008). Membranes were pre-hybridized in Ultrahyb Oligo hybridization buffer (Invitrogen), after which ^32^P labeled DNA oligonucleotides were added for overnight hybridization at 42°C. Membranes were washed 10 minutes in 2xSSC/0.1% SDS followed by 20 minutes in 1xSSC/0.1% SDS at 42°C. Kodak XAR X-ray films were exposed to radiolabeled membranes and developed using an Agfa CP1000 developer. Northern blots shown in Fig. 1 D, Fig. 7A and Supplemental Fig. S10C have already been used in Halbach et al. 2020, and were re-probed with a propiR1 probe. The miR2940-3p (Fig. 1D) and rRNA (Fig. 7A) images are shown here again as controls. Uncropped images are presented in Supplemental Fig. S11.

Probes used for northern blotting were:

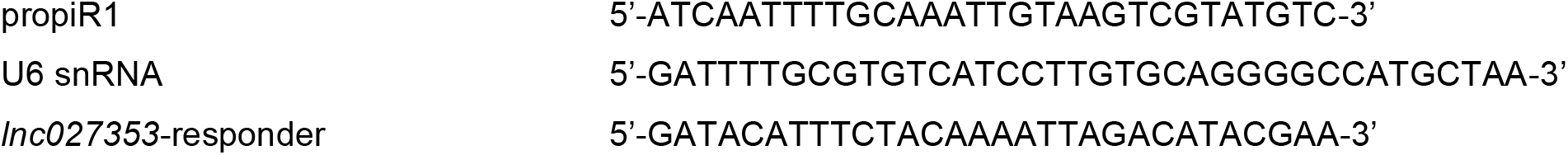

### Immunoprecipitation of PIWI proteins

Generation and characterization of anti-PIWI antibodies was described in Joosten et al. 2018 and Halbach et al. 2020 and immunoprecipitation was performed as described in Joosten et al. 2018. For Piwi4 and Piwi5 IP after knockdown of *Piwi5* and *Piwi4*, respectively, cells were subjected to three rounds of dsRNA mediated knockdown. The first dsRNA transfection was performed simultaneously with cell seeding, and the remainder of the knockdown procedure was performed as described above.

### Analysis of sequence conservation

Multiple sequence alignments (MSA) of DNA sequences were generated with ClustalW (https://www.genome.jp/tools-bin/clustalw) using default settings. Conservation scores for seven *Ae. aegypti* propiR1 sites plus 0.5 kb flanking regions were calculated using the consurf server (http://consurf.tau.ac.il/) (Ashkenazy et al. 2016) using the following settings: nucleotide sequence, no structure, MSA upload, no tree, Bayesian calculation method, Best evolutionary model. The conservation scores and the corresponding borders of the 50% confidence interval from the output file were multiplied by −1 to display higher conservation by increasing values and subsequently a sliding window analysis was applied. The mean of five consecutive nucleotide positions with an offset of one position was plotted.

### Reporter cloning

To generate reporter constructs, a pMT-GL3 vector, encoding firefly luciferase (GL3) under control of a copper sulphate (CuSO_4_)-inducible metallothionein promotor (pMT) (van Rij et al. 2006), was digested with *Pme*I and *Sac*II restriction enzymes for 3 to 4 hours at 37°C and dephosphorylated using Antarctic Phosphatase (NEB) for 1 hour at 37°C. To produce target site inserts, sense and antisense DNA oligonucleotides (50 μM) containing various propiR1 target sites (Supplemental Table S3) were heated at 90°C for 10 minutes in 100 mM Tris-HCl (pH 7.5), 0.1 M NaCl, 10 mM EDTA before gradually cooling to room temperature to anneal the oligonucleotides. Annealed oligonucleotides were subsequently phosphorylated for 30 minutes at 37°C using T4 polynucleotide kinase (Roche). 5 μL of 25 × diluted oligonucleotides and 50 ng of the digested and dephosphorylated vector were ligated overnight at 16°C using T4 Ligase (Roche), and transformed into XL10-Gold *E. coli*. Plasmid DNA was isolated using the High Pure Plasmid Isolation kit (Roche). Sequences were confirmed by Sanger sequencing.

### Luciferase assay

Approximately 2 × 10^4^ Aag2 cells/well were seeded in 96-well plates and incubated overnight. Per well, 100 ng of the pMT-GL3 reporter construct (van Rij et al. 2006) and 100 ng of pMT-*Renilla* Luciferase plasmid was transfected using 0.2 μL X-tremeGENE HP DNA, according to the manufacturer’s instructions. The pMT-*Renilla* Luciferase plasmid used in all experiments contained synonymous mutations in the tapiR1 target site to prevent Piwi4/tapiR1-mediated silencing (Halbach et al. 2020). To induce the metallothionein promotor of the reporter constructs, medium was replaced with fully supplemented Leibovitz’s L-15 medium containing 0.5 mM CuSO_4_ at 3 to 4 hours after transfection. In case of luciferase experiments in PIWI-depleted cells, reporter constructs were co-transfected with PIWI dsRNA during the second knockdown. In experiments in which PIWI-mediated silencing was blocked by antisense oligonucleotides (AOs), 100 ng of the FLuc reporters and 100 ng pMT-*Renilla* Luciferase were co-transfected with indicated concentrations of 5’ Cy5-labelled, fully 2’-*O*-methylated AOs using 4 μL X-tremeGENE HP DNA transfection reagent per μg of oligonucleotides. 24 hours after activation of the metallothionein promoters, cells were lysed in 30 μL Passive lysis buffer (Promega) per well. The activity of *Renilla* and firefly luciferase was measured using the Dual Luciferase Reporter Assay system (Promega) on a Modulus Single Tube Reader (Turner Biosystems). For each well, firefly luciferase was normalized to *Renilla* luciferase activity.

### mRNA-seq analysis

Reads were mapped to the *Aedes aegypti* LVP_AGWG AaegL5.1 reference genome obtained from VectorBase (https://www.vectorbase.org/) and quantified with STAR aligner (version 2.5.2b) (Dobin et al. 2013) in 2-pass mode. Briefly, all libraries were first mapped with options --readFilesCommand zcat --outSAMtype None --outSAMattrIHstart 448 0 --outSAMstrandField intronMotif, then all detected splice junctions were combined (false positive junctions on the mitochondrial genome were removed), and used in a second mapping step with –sjdbFileChrStartEnd, other parameters as above, and –quantMode GeneCounts in order to quantify gene expression. Transposons were quantified with Salmon (v.0.8.2) (Patro et al. 2017) on the TEfam transposon consensus sequence (https://tefam.biochem.vt.edu/tefam/, accessed April 2017) with default settings and libType set to “ISR”. Statistical analyses were performed with DEseq2 (Love et al. 2014) and significance was tested at an FDR of 0.01 and a log2 fold change of 0.5. Results were plotted in R with ggplot2 (Wickham 2009). Target sites for propiR1 on the differential regulated genes were predicted with the online tool of RNAhybrid available at https://bibiserv.cebitec.uni-bielefeld.de/rnahybrid/ (Rehmsmeier et al. 2004) with helix constraints from nt 2 to nt 7 of the piRNA and excluding G:U wobbles in this region.

For Supplemental Fig. S10A-B, publicly available datasets were obtained from Sequence Read Archive (SRA, https://www.ncbi.nlm.nih.gov/sra) and mapped and quantified with STAR aligner as described above. Normalization factors were estimated with DEseq2, and normalized counts were plotted with ggplot2. A list of datasets used can be found in Supplemental Table S4.

### sRNA-seq analysis

3’ sequencing adapters were clipped from sRNA sequencing reads with Cutadapt (version 1.14) (Martin 2011) (options -m 15 -M 35 --discard-untrimmed), that were subsequently mapped to the AaegL5.1 reference genome with Bowtie (version 0.12.17) (Langmead et al. 2009) without allowing mismatches (options --best -k 1 -v 0). Reads were trimmed to the first 25 nt to include different isoforms of a given piRNA, IP libraries (Miesen et al. 2015) were normalized to total mapped small RNA reads per million, and enrichment of piRNAs was then calculated as log2 fold enrichment compared to a GFP-control IP. For Fig. 1A, piRNAs that were at least twofold enriched in Piwi5 were filtered by their average abundance in triplicate control FLuc knockdown libraries from total small RNAs in Aag2 cells (Miesen et al. 2015), and overlapped with annotated transposable elements obtained from VectorBase with bedtools intersect (Quinlan 2014). Coverage of the *AAEL027353* target locus and the propiR1 loci were determined with bedtools coverage (Quinlan 2014), normalized to miRNA reads per million, and plotted in R with Gviz (Hahne & Ivanek 2016). Size distribution of propiR1 small RNAs in *Ae. aegypti* Aag2 and *Ae. albopictus* U4.4 cells were generated from the control knockdown condition from (Miesen et al. 2015) and from a small RNA sequencing library obtained from untreated U4.4 cells (PRJNA613255). Adapter sequences were clipped using the Clip adapter sequence tool (v1.03; options all default) within the public server of the usegalaxy.org instance (Afgan et al. 2018), and small RNA reads were subsequently mapped to the AaegL5.1 and the AalbF2 genome assemblies, respectively with Bowtie (version 0.12.17) (Langmead et al. 2009) without allowing mismatches (options: -n0 -l 32 -k 1). The read counts were normalized to the respective library size. To assess *lnc027353* targeting by Piwi5-associated piRNAs, all piRNA-sized sequences that were enriched at least 2-fold compared to a control GFP IP (Miesen et al. 2015) were extracted and used as input for target site prediction on the *lnc027353* in RNAhybrid, using the expressed part (regions covered by RNAseq reads) as input target sequence. In order to get an unbiased overview of piRNAs that are potentially able to target the lncRNA, the most favorable target site (lowest minimum free energy) was predicted for each piRNA without specifying any constraints. Average expression in Aag2 cells from three control FLuc knockdown libraries cells was plotted against the predicted minimum free energy, and all target sites that contained a seed match (nt 2-7) were highlighted. Nine target sites of the most abundant piRNAs (>500 rpm) were analyzed in a luciferase reporter assay.

### RT-qPCR

Cells were treated with AOs as described above. After RNA isolation, 0.5-1 μg of total RNA was DNaseI treated (Ambion), reverse transcribed using the Taqman reverse transcription kit (Applied Biosystems), and SYBR-green qPCR was performed using the GoTaq qPCR system (Promega) according to the manufacturers’ recommendations. Expression of target genes was normalized to the expression of the housekeeping gene lysosomal aspartic protease (LAP) and fold changes in expression were calculated using the 2^(-ΔΔCT)^ method (Livak & Schmittgen 2001).

### 5’ RACE of slicer products

To detect 3’ slicer products of propiR1-mediated cleavage events, a 5’ RACE RNA adapter was ligated to total Aag2 RNA with T4 RNA ligase (NEB) according to the manufacturer’s instructions. The reaction was reverse transcribed with Superscript IV (Invitrogen) using a transcript-specific reverse primer, and amplified with GoTaq Flexi polymerase (Promega) in two rounds of PCR (outer and inner), both with 5 min initial denaturation at 95°C, and 35 cycles of 30 s denaturation at 95°C, 30 s annealing at 60°C and 15 s extension at 72°C, followed by a final extension of 5 min. Homology arms complementary to the pUC19 cloning vector were then added to the PCR product with another PCR in order to clone the RACE products with the In-Fusion HD Cloning kit (Takara). Individual clones were selected and Sanger sequenced.

### Mosquito rearing and egg laying

*Ae. aegypti* mosquitoes Liverpool strain were reared at 28±1 °C, 80% humidity with a 12h:12h light:dark cycle. Eggs were hatched in tap water, and larvae were fed with fish food powder (Tetramin) every other day. Adults were allowed constant access to 10% (w/v) sucrose in water, and females were fed on human blood (Sanquin Blood Supply Foundation, Nijmegen, the Netherlands) through a membrane feeding system (Hemotek Ltd.). For injection experiments, females were separated, offered a blood meal, and allowed to lay eggs three to four days later by providing a moist surface and placing the mosquitoes in the dark. Embryos were collected at the indicated time points after a 30-minute egg laying period. For Fig. 7A-B, *Ae. aegypti* Jane mosquitoes were used as described previously (Halbach et al. 2020).

### Embryo injections

Pre-blastoderm embryos were injected as described previously (Aryan et al. 2014). Briefly, gravid females were allowed to lay eggs for 45 min, then embryos were aged for 1-2 hours until they turned gray, desiccated for 1.5 min, covered with halocarbon oil and injected with 50 μM antisense oligonucleotides with a Pneumatic PicoPump PV820 with 30 psi inject pressure. Injected embryos were then transferred to a wet Whatman paper, kept at 28°C and 80% humidity and harvested 20 hours later. For each condition, ~50 eggs were injected and pooled prior to RNA isolation.

### Statistical analyses

Unless noted otherwise, unpaired two tailed t-tests with Holm-Sidak correction for multiple comparisons were used to test statistical significance (* *P* < 0.05, ** *P* < 0.005, *** *P* < 0.0005).

### Data availability

RNA sequencing of propiR1 AO-treated *Aedes aegypti* Aag2 cells and small RNA libraries from *Aedes albopictus* U4.4 cells were deposited in the NCBI Sequence Read Archive under BioProject numbers PRJNA622708 and PRJNA613255, respectively.

## ACKNOWLEDGEMENTS

We thank members of the laboratory for fruitful discussions, Bas Pennings for technical assistance in cloning of luciferase reporters, and Prof. Martijn Huijnen from the Center for Molecular and Biomolecular Informatics (CMBI), Nijmegen, for bioinformatics advice. Sequencing was performed by the GenomEast platform, a member of the “France Génomique” consortium (ANR-10-INBS-0009). This work was financially supported by a fellowship from Radboud University Medical Center, a Consolidator Grant from the European Research Council (ERC) under the European Union’s Seventh Framework Programme (ERC CoG grant number 615680), and a VICI grant from the Dutch Research Council (NWO, grant number 016.VICI.170.090). *Ae. aegypti* Liverpool strain mosquitoes were kindly provided by the NIH/NIAID Filariasis Research Reagent Resource Center, distributed by BEI Resources, NIAID, NIH.

## REFERENCE LIST

Afgan E, Baker D, Batut B, van den Beek M, Bouvier D, Cech M, Chilton J, Clements D, Coraor N, Gruning BA, et al. 2018. The Galaxy platform for accessible, reproducible and collaborative biomedical analyses: 2018 update. Nucleic acids research 46: W537–W544

Akbari OS, Antoshechkin I, Amrhein H, Williams B, Diloreto R, Sandler J, Hay BA. 2013. The developmental transcriptome of the mosquito Aedes aegypti, an invasive species and major arbovirus vector. G3 (Bethesda) 3: 1493–509

Arensburger P, Hice RH, Wright JA, Craig NL, Atkinson PW. 2011. The mosquito Aedes aegypti has a large genome size and high transposable element load but contains a low proportion of transposon-specific piRNAs. BMC Genomics 12: 606

Aryan A, Myles KM, Adelman ZN. 2014. Targeted genome editing in Aedes aegypti using TALENs. Methods 69: 38–45

Ashkenazy H, Abadi S, Martz E, Chay O, Mayrose I, Pupko T, Ben-Tal N. 2016. ConSurf 2016: an improved methodology to estimate and visualize evolutionary conservation in macromolecules. Nucleic acids research 44: W344–50

Barckmann B, Pierson S, Dufourt J, Papin C, Armenise C, Port F, Grentzinger T, Chambeyron S, Baronian G, Desvignes J-P, et al. 2015. Aubergine iCLIP Reveals piRNA-Dependent Decay of mRNAs Involved in Germ Cell Development in the Early Embryo. Cell Reports 12: 1205–1216

Brennecke J, Aravin AA, Stark A, Dus M, Kellis M, Sachidanandam R, Hannon GJ. 2007. Discrete small RNA-generating loci as master regulators of transposon activity in Drosophila. Cell 128: 1089–103

Brennecke J, Malone CD, Aravin AA, Sachidanandam R, Stark A, Hannon GJ. 2008. An epigenetic role for maternally inherited piRNAs in transposon silencing. Science (New York, NY) 322: 1387–1392

Campbell CL, Black WC, Hess AM, Foy BD. 2008. Comparative genomics of small RNA regulatory pathway components in vector mosquitoes. BMC Genomics 9: 425

Crava CM, Varghese FS, Pischedda E, Halbach R, Palatini U, Marconcini M, Gasmi L, Redmond S, Afrane Y, Ayala D, et al. 2021. Population genomics in the arboviral vector Aedes aegypti reveals the genomic architecture and evolution of endogenous viral elements. Molecular Ecology, 30(7), 1594–1611

Czech B, Hannon GJ. 2016. One Loop to Rule Them All: The Ping-Pong Cycle and piRNA-Guided Silencing. Trends Biochem Sci 41: 324–337

Danet L, Beauclair G, Berthet M, Moratorio G, Gracias S, Tangy F, Choumet V, Jouvenet N. 2019. Midgut barriers prevent the replication and dissemination of the yellow fever vaccine in Aedes aegypti. PLoS Negl Trop Dis 13: e0007299

Dobin A, Davis CA, Schlesinger F, Drenkow J, Zaleski C, Jha S, Batut P, Chaisson M, Gingeras TR. 2013. STAR: ultrafast universal RNA-seq aligner. Bioinformatics 29: 15–21

Girardi E, Miesen P, Pennings B, Frangeul L, Saleh MC, van Rij RP. 2017. Histone-derived piRNA biogenesis depends on the ping-pong partners Piwi5 and Ago3 in Aedes aegypti. Nucleic Acids Res 45: 4881–4892

Gunawardane LS, Saito K, Nishida KM, Miyoshi K, Kawamura Y, Nagami T, Siomi H, Siomi MC. 2007. A slicer-mediated mechanism for repeat-associated siRNA 5’ end formation in Drosophila. Science 315: 1587–90

Hahne F, Ivanek R. 2016. Visualizing Genomic Data Using Gviz and Bioconductor. Methods in molecular biology 1418: 335–351

Halbach R, Miesen P, Joosten J, Taşköprü E, Rondeel I, Pennings B, Vogels CBF, Merkling SH, Koenraadt CJ, Lambrechts L, et al. 2020. A satellite repeat-derived piRNA controls embryonic development of Aedes. Nature 580: 274–277

Han BW, Wang W, Li CJ, Weng ZP, Zamore PD. 2015. piRNA-guided transposon cleavage initiates Zucchini-dependent, phased piRNA production. Science 348: 817–821

Horwich MD, Li C, Matranga C, Vagin V, Farley G, Wang P, Zamore PD. 2007. The Drosophila RNA methyltransferase, DmHen1, modifies germline piRNAs and single-stranded siRNAs in RISC. Current biology 17: 1265–72

Jensen S, Brasset E, Parey E, Roest Crollius H, Sharakhov IV, Vaury C. 2020. Conserved Small Nucleotidic Elements at the Origin of Concerted piRNA Biogenesis from Genes and lncRNAs. Cells 9

Joosten J, Miesen P, Taşköprü E, Pennings B, Jansen PWTC, Huynen MA, Vermeulen M, Van Rij RP. 2018. The Tudor protein Veneno assembles the ping-pong amplification complex that produces viral piRNAs in Aedes mosquitoes. Nucleic Acids Research 47: 2546–2559

Joosten J, Van Rij RP, Miesen P. 2020. Slicing of viral RNA guided by endogenous piRNAs triggers the production of responder and trailer piRNAs in Aedes mosquitoes. bioRxiv doi:10.1101/2020.07.08.193029

Kawaoka S, Katsuma S, Tomari Y. 2014. Making piRNAs in vitro. Methods in molecular biology 1093: 35–46

Kobayashi H, Tomari Y. 2016. RISC assembly: Coordination between small RNAs and Argonaute proteins. Bba-Gene Regul Mech 1859: 71–81

Langmead B, Trapnell C, Pop M, Salzberg SL. 2009. Ultrafast and memory-efficient alignment of short DNA sequences to the human genome. Genome biology 10: R25

Lewis SH, Quarles KA, Yang YJ, Tanguy M, Frezal L, Smith SA, Sharma PP, Cordaux R, Gilbert C, Giraud I, et al. 2018. Pan-arthropod analysis reveals somatic piRNAs as an ancestral defence against transposable elements. Nat Ecol Evol 2: 174–181

Lewis SH, Salmela H, Obbard DJ. 2016. Duplication and Diversification of Dipteran Argonaute Genes, and the Evolutionary Divergence of Piwi and Aubergine. Genome Biol Evol 8: 507–18

Liu P, Dong Y, Gu J, Puthiyakunnon S, Wu Y, Chen XG. 2016. Developmental piRNA profiles of the invasive vector mosquito Aedes albopictus. Parasites & vectors 9: 524

Livak KJ, Schmittgen TD. 2001. Analysis of relative gene expression data using real-time quantitative PCR and the 2(-Delta Delta C(T)) Method. Methods 25: 402–8

Love MI, Huber W, Anders S. 2014. Moderated estimation of fold change and dispersion for RNA-seq data with DESeq2. Genome biology 15: 550

Martin M. 2011. Cutadapt removes adapter sequences from high-throughput sequencing reads. EMBnetjournal 17: pg. 10–12

Matsumoto N, Nishimasu H, Sakakibara K, Nishida KM, Hirano T, Ishitani R, Siomi H, Siomi MC, Nureki O (2016) Crystal Structure of Silkworm PIWI-Clade Argonaute Siwi Bound to piRNA. Cell 167: 484–497 e9

Matthews BJ, McBride CS, DeGennaro M, Despo O, Vosshall LB. 2016. The neurotranscriptome of the Aedes aegypti mosquito. Bmc Genomics 17: 32

Miesen P, Girardi E, van Rij RP. 2015. Distinct sets of PIWI proteins produce arbovirus and transposon-derived piRNAs in Aedes aegypti mosquito cells. Nucleic Acids Res 43: 6545–56

Miesen P, Joosten J, van Rij RP. 2016. PIWIs Go Viral: Arbovirus-Derived piRNAs in Vector Mosquitoes. PLoS pathogens 12: e1006017–e1006017

Mohn F, Handler D, Brennecke J. 2015. piRNA-guided slicing specifies transcripts for Zucchini-dependent, phased piRNA biogenesis. Science 348: 812–817

Morazzani EM, Wiley MR, Murreddu MG, Adelman ZN, Myles KM. 2012. Production of Virus-Derived Ping-Pong-Dependent piRNA-like Small RNAs in the Mosquito Soma. PLOS Pathogens 8: e1002470

Ozata DM, Gainetdinov I, Zoch A, O’Carroll D, Zamore PD. 2019. PIWI-interacting RNAs: small RNAs with big functions. Nat Rev Genet 20: 89–108

Palatini U, Miesen P, Carballar-Lejarazu R, Ometto L, Rizzo E, Tu Z, van Rij RP, Bonizzoni M. 2017. Comparative genomics shows that viral integrations are abundant and express piRNAs in the arboviral vectors Aedes aegypti and Aedes albopictus. BMC Genomics 18: 512

Pall GS, Hamilton AJ. 2008. Improved northern blot method for enhanced detection of small RNA. Nature Protocols 3: 1077–1084

Patro R, Duggal G, Love MI, Irizarry RA, Kingsford C. 2017. Salmon provides fast and bias-aware quantification of transcript expression. Nature methods 14: 417–419

Quinlan AR. 2014. BEDTools: The Swiss-Army Tool for Genome Feature Analysis. Current protocols in bioinformatics 47: 11 12 1–34

Rehmsmeier M, Steffen P, Hochsmann M, Giegerich R. 2004. Fast and effective prediction of microRNA/target duplexes. Rna 10: 1507–17

Reuter JS, Mathews DH. 2010. RNAstructure: software for RNA secondary structure prediction and analysis. BMC Bioinformatics 11: 129–129

Reuter M, Berninger P, Chuma S, Shah H, Hosokawa M, Funaya C, Antony C, Sachidanandam R, Pillai RS. 2011. Miwi catalysis is required for piRNA amplification-independent LINE1 transposon silencing. Nature 480: 264–7

Rouget C, Papin C, Boureux A, Meunier AC, Franco B, Robine N, Lai EC, Pelisson A, Simonelig M. 2010. Maternal mRNA deadenylation and decay by the piRNA pathway in the early Drosophila embryo. Nature 467: 1128–32

Saito K, Sakaguchi Y, Suzuki T, Suzuki T, Siomi H, Siomi MC. 2007. Pimet, the Drosophila homolog of HEN1, mediates 2’-O-methylation of Piwi-interacting RNAs at their 3’ ends. Genes & development 21: 1603–8

Schnettler E, Donald CL, Human S, Watson M, Siu RWC, McFarlane M, Fazakerley JK, Kohl A, Fragkoudis R. 2013. Knockdown of piRNA pathway proteins results in enhanced Semliki Forest virus production in mosquito cells. J Gen Virol 94: 1680–1689

Shen EZ, Chen H, Ozturk AR, Tu S, Shirayama M, Tang W, Ding YH, Dai SY, Weng Z, Mello CC. 2018. Identification of piRNA Binding Sites Reveals the Argonaute Regulatory Landscape of the C. elegans Germline. Cell 172: 937–951.e18

Suzuki Y, Frangeul L, Dickson LB, Blanc H, Verdier Y, Vinh J, Lambrechts L, Saleh MC. 2017. Uncovering the Repertoire of Endogenous Flaviviral Elements in Aedes Mosquito Genomes. J Virol 91

Tadros W, Lipshitz HD. 2009. The maternal-to-zygotic transition: a play in two acts. Development 136: 3033–42

Tassetto M, Kunitomi M, Whitfield ZJ, Dolan PT, Sánchez-Vargas I, Garcia-Knight M, Ribiero I, Chen T, Olson KE, Andino R. 2019. Control of RNA viruses in mosquito cells through the acquisition of vDNA and endogenous viral elements. eLife 8: e41244

Tian Y, Simanshu DK, Ma JB, Patel DJ. 2011. Structural basis for piRNA 2’-O-methylated 3’-end recognition by Piwi PAZ (Piwi/Argonaute/Zwille) domains. Proceedings of the National Academy of Sciences of the United States of America 108: 903–10

van Rij RP, Saleh MC, Berry B, Foo C, Houk A, Antoniewski C, Andino R. 2006. The RNA silencing endonuclease Argonaute 2 mediates specific antiviral immunity in Drosophila melanogaster. Genes & development 20: 2985–2995

Vodovar N, Bronkhorst AW, van Cleef KWR, Miesen P, Blanc H, van Rij RP, Saleh MC. 2012. Arbovirus-Derived piRNAs Exhibit a Ping-Pong Signature in Mosquito Cells. PLOS ONE 7: e30861

Wang Y, Jin B, Liu P, Li J, Chen X, Gu J. 2018. piRNA Profiling of Dengue Virus Type 2-Infected Asian Tiger Mosquito and Midgut Tissues. Viruses 10

Whitfield ZJ, Dolan PT, Kunitomi M, Tassetto M, Seetin MG, Oh S, Heiner C, Paxinos E, Andino R. 2017. The Diversity, Structure, and Function of Heritable Adaptive Immunity Sequences in the Aedes aegypti Genome. Curr Biol 27: 3511–3519.e7

Wickham H. 2009. Ggplot2 : elegant graphics for data analysis. Springer, New York

Yamaguchi S, Oe A, Nishida KM, Yamashita K, Kajiya A, Hirano S, Matsumoto N, Dohmae N, Ishitani R, Saito K, et al. 2020. Crystal structure of Drosophila Piwi. Nat Commun 11: 858

