## Supplemental Figures S1 to S11 for "A piRNA-lncRNA regulatory network initiates responder and trailer piRNA formation during mosquito embryonic development"

### *Supplemental Tables*

Supplemental Table S1 – Differentially expressed genes in Aag2 cells treated with propiR1 antisense oligonucleotides.

Supplemental Table S2 – Differentially expressed transposons in Aag2 cells treated with propiR1 antisense oligonucleotides.

Supplemental Table S3 – Oligonucleotide sequences

Supplemental Table S4 – RNAseq datasets analyzed in this study

### *Supplemental Figures*

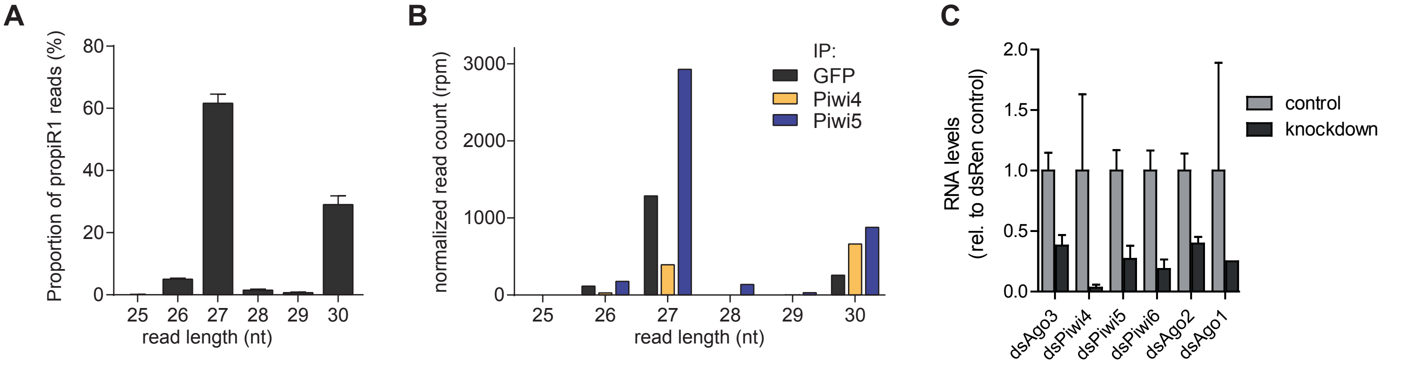


**Supplemental Fig. S1. Size distribution of propiR1 isoforms in Ae. aegypti.** A) Size distribution of propiR1 reads in small RNA deep sequencing libraries generated from Aag2 cells treated with dsRNA targeting luciferase. B) Size distribution of propiR1 isoforms in immunoprecipitations (IP) of V5 epitope-tagged Piwi4 and Piwi5. V5-GFP-IP serves as non-specific background binding control. Read count is normalized to library size. The datasets used for (A) and (B) were first described in (Miesen et al., 2015). C) Knockdown efficiencies in Aag2 cells transfected with dsRNA targeting the indicated genes measured by RT-qPCR, normalized to control dsRNA (targeting *Renilla* luciferase). The same RNA was used in Figure 1E. Bars and whiskers depict the mean +/- SD of three biological replicates.

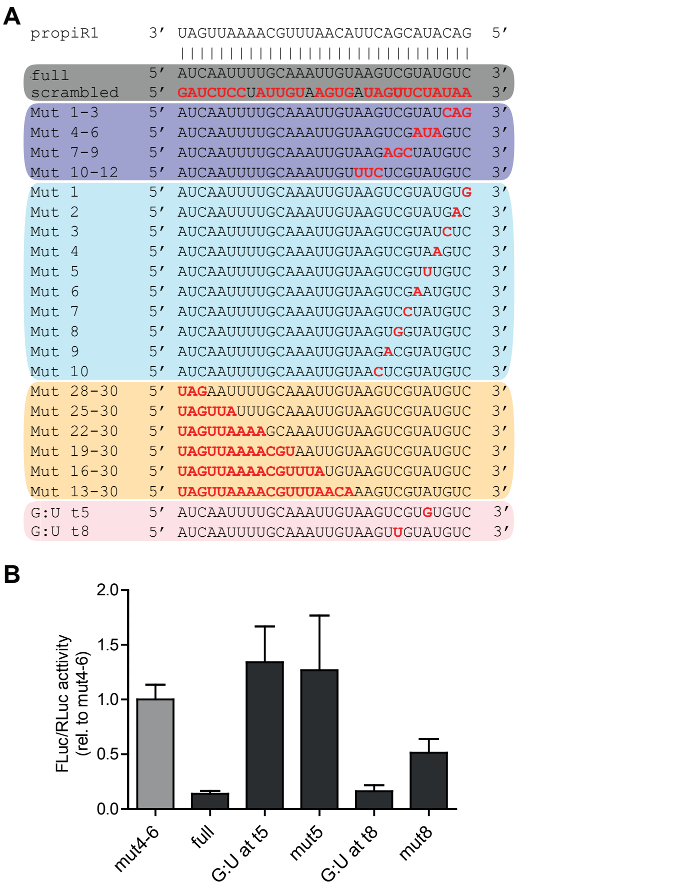


**Supplemental Fig. S2. Target site mutations in propiR1 luciferase reporters.** A) Schematic overview of mutations introduced into the propiR1 target site, used to study targeting requirements. Gray shaded area includes the positive (fully complementary) and negative (scrambled) target site controls. Dark blue, light blue and yellow shaded areas indicate triplet seed mutants, single nt seed mutants, and 3’ region mutants, respectively. Colors correspond to those used in Figure 2C. B) Luciferase assay of reporters bearing G:U wobble base pairs in the propiR1 seed-target RNA duplex. A propiR1 target site in which residues t4-6 were mutated (mut4-6) serves as control. Bars represent the mean +/- SD of a representative of two independent experiments, each performed with three biological replicates.

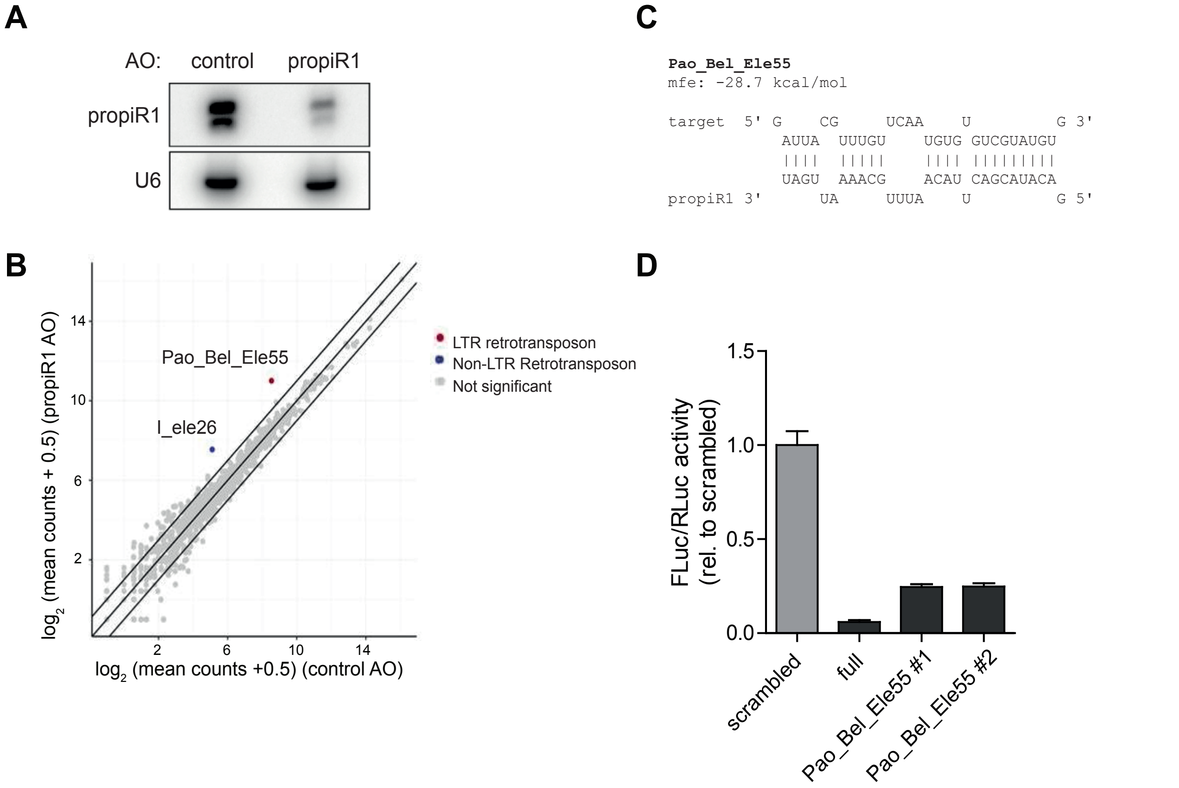


**Supplemental Fig. S3. Effect of propiR1 on RNA expression of transposable elements.** A) Northern blot analysis of propiR1 in Aag2 cells treated with 300 nM propiR1 or control antisense oligonucleotides (AO) for 48 hours. B) Log2 expression of transposable elements in Aag2 cells treated with propiR1 or control AOs. Mean RNA-seq counts of three biological replicates are shown (plus a pseudo-count of 0.5 to plot values of zero). Significance was tested at an FDR of 0.01 and log2 fold change of 0.5. Diagonal lines highlight a 2-fold change. Two transposons were differentially expressed, *Pao_Bel_Ele55* and *I_ele26*. Of these two, only *Pao_Bel_Ele55* had a seed-based target site predicted by RNAhybrid. C) Schematic representation of predicted propiR1-target duplex in *Pao_Bel_Ele55*. D) Luciferase assay of a reporter bearing the predicted target site in *Pao_Bel_Ele55* . Data are normalized to the scrambled control reporter (scrambled). Bars represent mean +/- SD of three biological replicates of a representative of two independent experiments. Suffixes 1 and 2 indicate two different clones.

**A**

```
AAEL027353-RA
mfe: -29.4 kcal/mol

target  5' U    GC        GGCAAU         U 3'
            UCGG  UUGCAGAU      UCGUAUGUC
            ||||  ||||||||      |||||||||
            AGUU  AACGUUUA      AGCAUACAG
propiR1 3' U    AA        ACAUUC           5

AAEL001794-RB
mfe: -23.6 kcal/mol

target  5'  A          CUCACCG           C 3'
             CGGUUUUGCG          CGUAUGUU
             ||||||||||          ||||||||
             GUUAAAACGU          GCAUACAG
propiR1 3' UA          UUAACAUUCA          5'

AAEL000564-RH
mfe: -26.3 kcal/mol

target  5'         A    CAUCAAU            A 3'
                    GCAG       GGUCGUAUGUC
                    ||||       |||||||||||
                    CGUU       UCAGCAUACAG
propiR1 3' UAGUUAAAA    UAACAU               5'

AAEL006445-RA
mfe: -24.2 kcal/mol

target  5' U   GG    G CUGAUG   GUUGGG     A         U 3'
            UCG  UUUG G      GAU      GUGGG UUGUAUGUC
            |||  |||| |      |||      ||||| |||||||||
            AGU  AAAC U      UUA      CAUUC AGCAUACAG
propiR1 3' U   UA    G         A                       5'

AAEL003865-RB
mfe: -24.4 kcal/mol

target 5'  G    CGA        A  AUCG   GAA       A 3'
            AUCA     GCAGGU GU    GAG   GUAUGUC
            ||||     |||||| ||    |||   |||||||
            UAGU     CGUUUA CA    UUC   CAUACAG
propiR1 3'      UAAAA      A         AG          5'
```

**B**

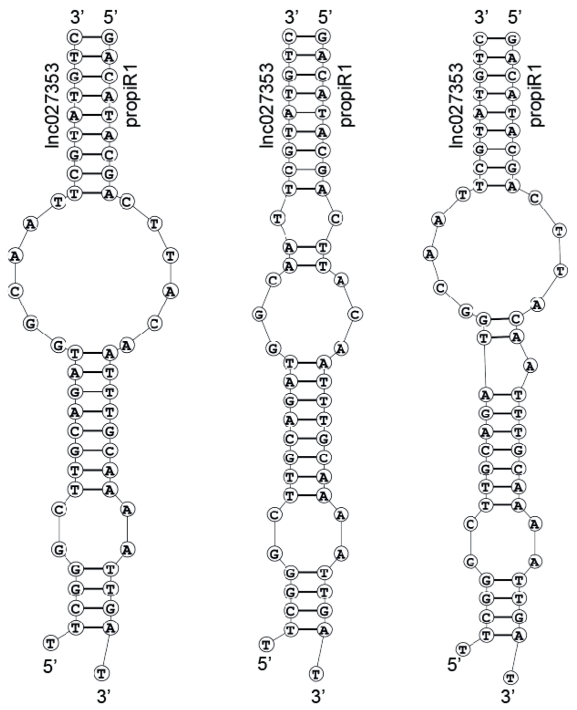


**Supplemental Fig. S4. Predicted structures of propiR1-target duplexes.** A) Schematic representation of predicted propiR1-target duplexes (by RNAhybrid) in the indicated transcripts and their minimum free energy (mfe). B) Alternative structures predicted by Bifold. The leftmost structure matches the prediction by RNAhybrid that was used in the manuscript.

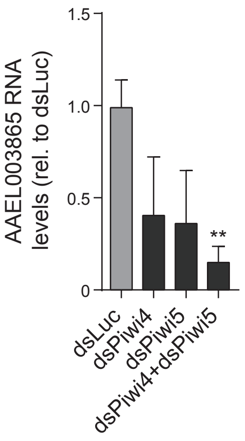


**Supplemental Fig. S5. Expression of *AAEL003685* upon Piwi4 and Piwi5 knockdown.** Relative expression of *AAEL003685* upon *Piwi4* and *Piwi5* single and double knockdown in Aag2 cells, measured by RT-qPCR. Cells treated with dsRNA targeting Luciferase (dsLuc) serves as a control. *Piwi4* and *Piwi5* knockdown resulted in decreased *AAEL003865* expression (~2.4-fold and ~2.7-fold, respectively) and combined *Piwi4* and *Piwi5* knockdown resulted in a ~6.3-fold reduction of *AAEL003865* expression. These results are in line with the finding that *AAEL003685* expression was reduced upon propiR1 AO treatment (Figure 3B-C). Yet, as the propiR1 target site predicted in *AAEL003685* did not affect luciferase activity in reporter assays (Figure 3E), propiR1 is unlikely to directly regulate *AAEL003865* expression. Bars represent the mean +/- SD of three biological replicates. Asterisks denote statistically significant differences in *AAEL003685* expression compared to dsLuc treated cells (unpaired two tailed t-tests with Holm-Sidak correction; ** $P < 0.005$).

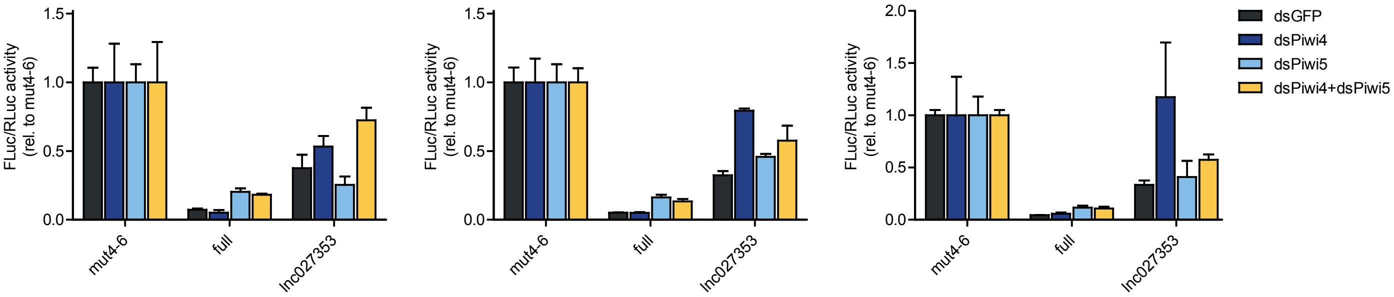


**Supplemental Fig. S6. Silencing of reporters with the endogenous target site of *lnc027353* upon PIWI knockdown.** Luciferase assays of reporters bearing a fully complementary propiR1 target site (full), or the endogenous target site of *lnc027353* after single or double knockdown of *Piwi4* and *Piwi5*. A propiR1 target site in which residues t4-6 were mutated (mut4-6) serves as control. Luciferase activity was normalized to the mut4-6 control, treated with the same dsRNA. Three independent experiments are shown, each performed with three biological replicates. Bars represent mean +/- SD.

A

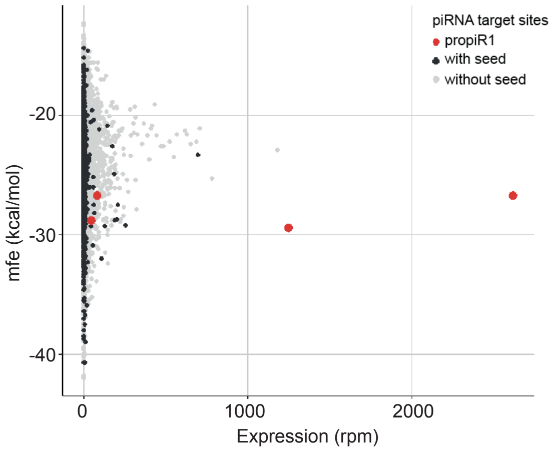


B

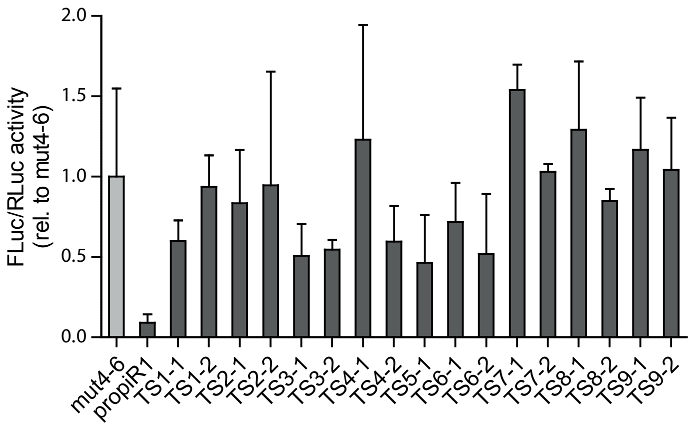


**Supplemental Fig. S7. *lnc027353* is not targeted by other Piwi5-associated piRNAs.** A) Dot plot of all Piwi5-associated piRNAs with a predicted target site in *lnc027353*, based on their expression and the minimal free energy (mfe) of the predicted propiR1:target duplex. Red dots indicate propiR1 isoforms of different lengths. B) Luciferase assay of reporters with target sites for the nine most abundant piRNAs (expression > 500 rpm) predicted to target *lnc027353* at non-propiR1 overlapping sites. The propiR1 target site in *lnc027353* was included as a positive control. Bars represent the mean +/- SD of three biological replicates, from a representative of two independent experiments. Suffixes -1 and -2 refer to two independent clones for each reporter that have been tested.

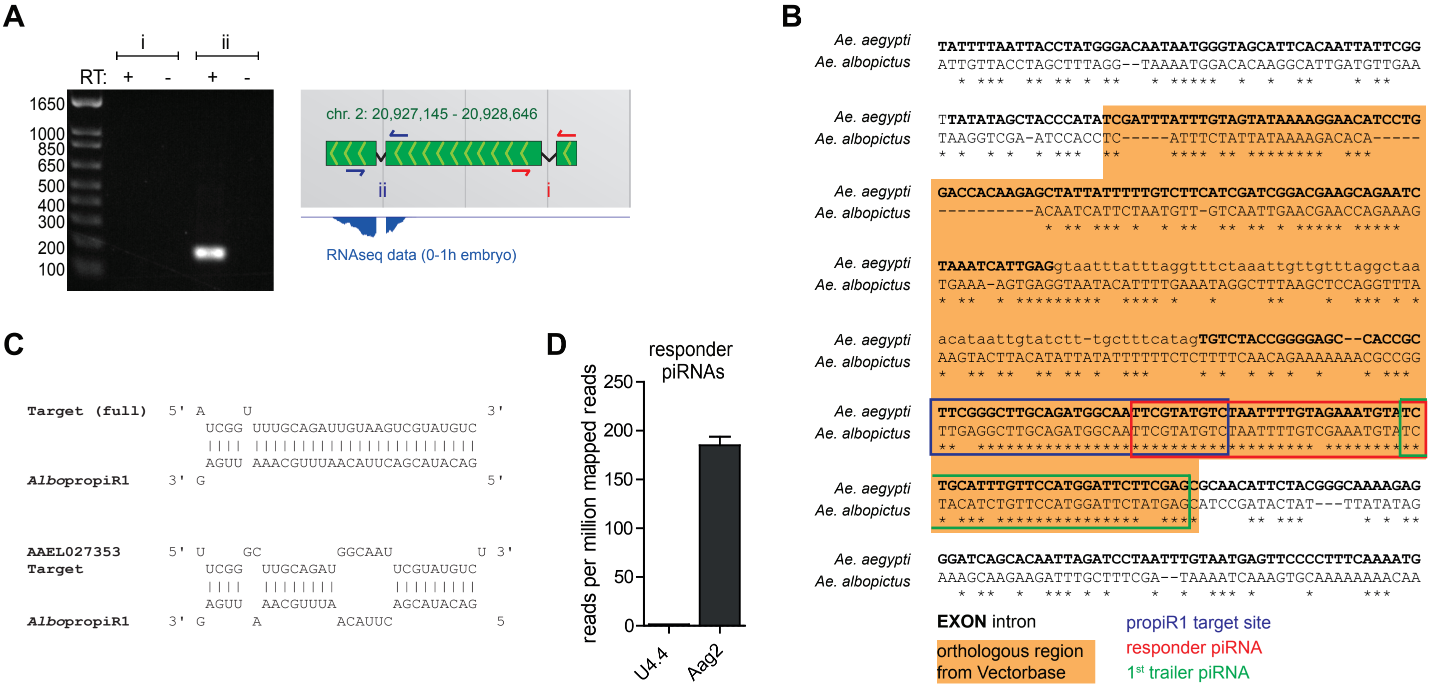


**Supplemental Fig. S8. Characterization of the *lnc027353* orthologous sequence in *Ae. albopictus*.** A) PCR on cDNA from *Ae. aegypti* Aag2 cells with primer pairs covering either the exon 1-2 junction (i) or the exon 2-3 junction (ii) (primer location shown in the right panel). Reactions in the absence of reverse transcriptase (RT) were included as control, as indicated. B) Alignment of *Ae. aegypti lnc027353* and its *Ae. albopictus* ortholog *Aalb_lnc027353.* The shaded area indicates a stretch that is annotated in VectorBase as an orthologous region. The propiR1 target site, responder piRNA, and the 1st trailer piRNA are indicated by blue, red and green boxes, respectively. For *Ae. aegypti*, sequences annotated as exons are in uppercase bold font; intronic sequence is in lowercase regular font. C) Schematic representation of piRNA-target RNA duplexes of *Ae. albopictus* propiR1 with the *Ae. aegypti* propiR1 fully complementary target site (upper panel) and the *Ae. aegypti lnc027353* target site (lower panel) in the luciferase reporters used in Figure 6D. D) Number of responder piRNAs per million mapped reads (rpm) that are generated from the *lnc027353* transcript in *Ae. aegypti* Aag2 cells and its ortholog in *Ae. albopictus* U4.4 cells. Only piRNAs with an exact 10 nt 5'-5' offset to propiR1 are plotted. Indicated are mean +/- SD from triplicate libraries for Aag2 cells and the value of a single library for U4.4 cells.

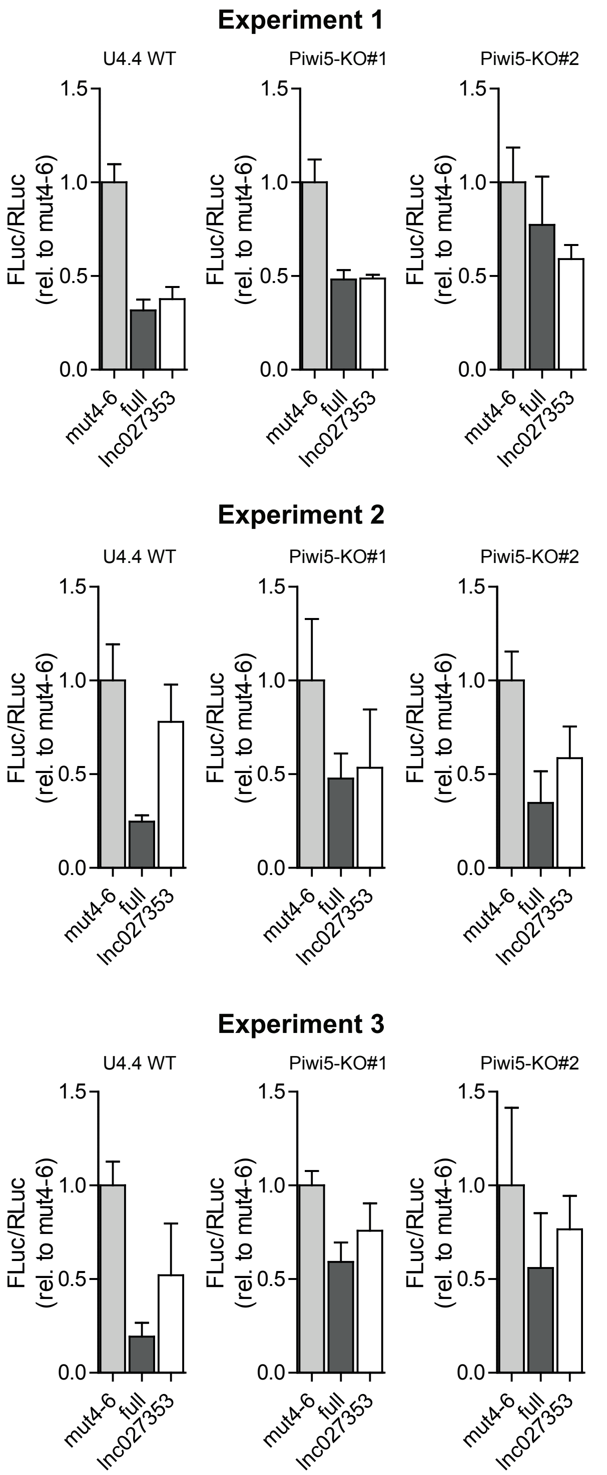


**Supplemental Fig. S9. Silencing of reporters with the endogenous target site of *lnc027353* in U4.4 WT and Piwi5 KO cells.** Luciferase assay in WT and *Piwi5* knockout (KO) U4.4 cells using reporters containing a control target site with mismatches at positions t4-6 (mut4-6), a fully complementary propiR1 target site (full) or the endogenous target site from *Ae. aegypti lnc027353*. Data were normalized to the activity of a co-transfected *Renilla* luciferase reporter (RLuc) and expression relative to mut4-6 is shown. Shown are the three independent replicate experiments of Figure 6D. Bars depict mean +/- SD of three biological replicates.

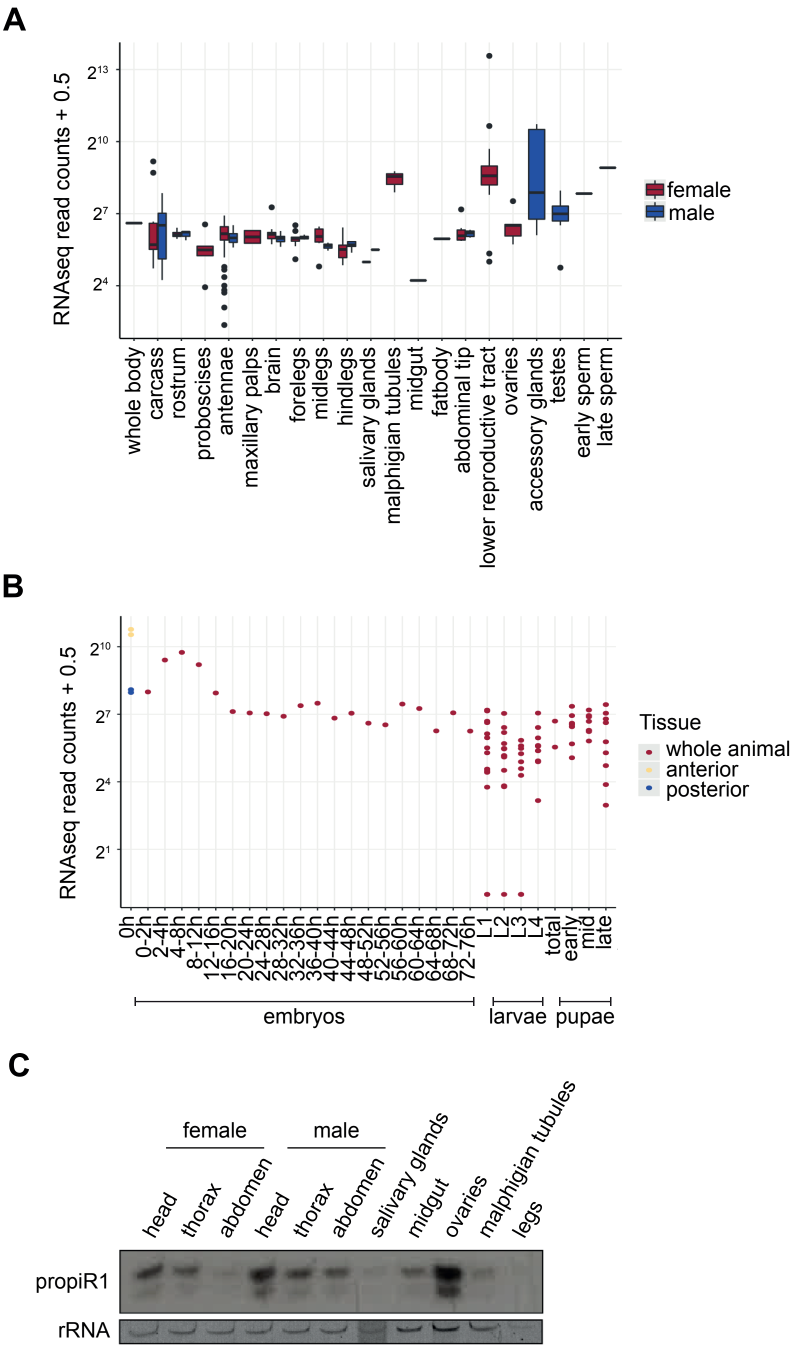


**Supplemental Fig. S10. Spatiotemporal expression of *lnc027353* and propiR1.** A-B) Expression of *lnc027353* in different tissues (A) and across developmental stages (B) of *Ae. aegypti.* Values are normalized RNA-seq counts with a pseudo-count of 0.5 added to plot values of zero. Shown are box-whisker plots with median, 1st and 3rd quantile and outlying points (A), and read counts in individual libraries (B). C) Northern blot analysis of propiR1 in different tissues of adult *Ae. aegypti.* EtBr stained rRNA serves as a loading control. The northern blot was published in (Halbach et al., 2020), and re-probed for propiR1.

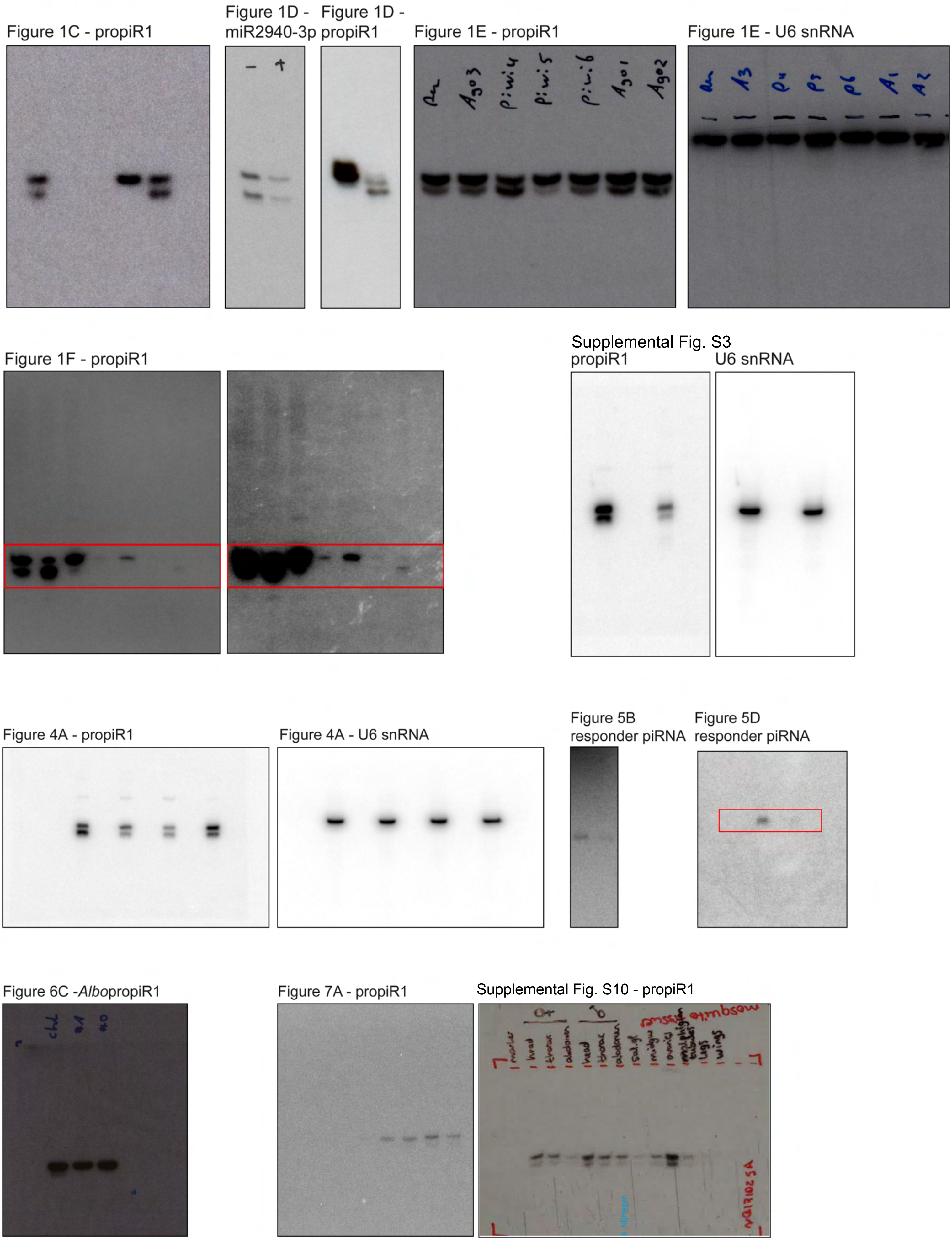


**Supplemental Fig. S11 - Uncropped images**
